# Sex differences in neural activity across amygdalo-striatal network during social behaviour

**DOI:** 10.64898/2026.05.25.727611

**Authors:** Adèle Phalip, Shai Netser, Shlomo Wagner

## Abstract

**Background:** Social behaviour is essential for the survival of most mammalian species and is shaped by sex-dependent genetic and endocrine factors. However, how sex influences brain-wide neural dynamics during social interactions remains poorly understood.

**Methods:** Here, we investigated sex differences in neural activity across a distributed amygdalo-striatal network in freely behaving mice. Using chronically implanted electrode arrays, we simultaneously recorded extracellular activity from multiple amygdalo-striatal regions while mice performed four social discrimination tasks. Neural signals were analysed alongside video-based behavioural tracking and head acceleration measurements.

**Results:** We identified significant sex differences in neural activity that emerged even before social interaction, suggesting distinct anticipatory network states. During social interaction, sex differences were distributed across brain regions and electrophysiological features, but were most consistently expressed in the basolateral amygdala (BLA). Notably, BLA activity exhibited pronounced sex-specific, context- and time-dependent dynamics, particularly during the initial phase of social interaction. These neural differences were associated with variations in behavioural responses and movement dynamics.

**Conclusions:** Together, our findings reveal that sex shapes both baseline and interaction-driven neural activity across the social brain network, and highlight the BLA as a key node underlying sex-specific dynamics of social behaviour.

**Plain English summary:** Social behaviour is essential for survival and differs between males and females in many species, including humans and mice. These differences are influenced by biological factors such as genes and hormones, but how they are reflected in brain activity during social interactions is still not fully understood.

In this study, we examined how brain activity differs between male and female mice during social behaviour. We recorded neural activity simultaneously from several brain regions involved in social and emotional processing while mice performed four different social interaction tasks. These tasks tested preferences for social versus non-social stimuli, opposite-sex animals, isolated animals, and stressed animals. At the same time, we monitored behaviour and head movements using video tracking and motion sensors.

We found that males and females showed distinct patterns of brain activity even before social interaction began, suggesting that the brain may enter different “anticipatory” states depending on sex. During social interaction, sex differences in neural activity varied depending on the social context and the stage of the interaction. The strongest and most consistent differences were observed in the basolateral amygdala, a brain region known to regulate emotional and social behaviour.

Interestingly, these neural differences were linked to differences in movement dynamics and social responses, particularly during the first moments of interaction. Our findings suggest that sex shapes both baseline brain activity and the way the brain responds during social encounters. This work improves our understanding of the neural basis of sex differences in social behaviour and may help inform future research on psychiatric conditions that affect social functioning differently in males and females.

**Highllights:**

- Simultaneous multi-site recordings revealed sex-dependent neural dynamics across an amygdalo-striatal social brain network during social behaviour.
- Male and female mice exhibited distinct electrophysiological signatures even before social interaction, suggesting sex-specific anticipatory neural states.
- High-frequency local field potential oscillations showed the strongest and most consistent sex differences across behavioural contexts and brain regions.
- The basolateral amygdala (BLA) emerged as a key region displaying context- and time-dependent sex differences during early social interaction.
- Sex-specific BLA activity correlated with movement dynamics during investigation of isolated conspecifics, linking neural network activity to behavioural responses.

**Graphical Abstract:** 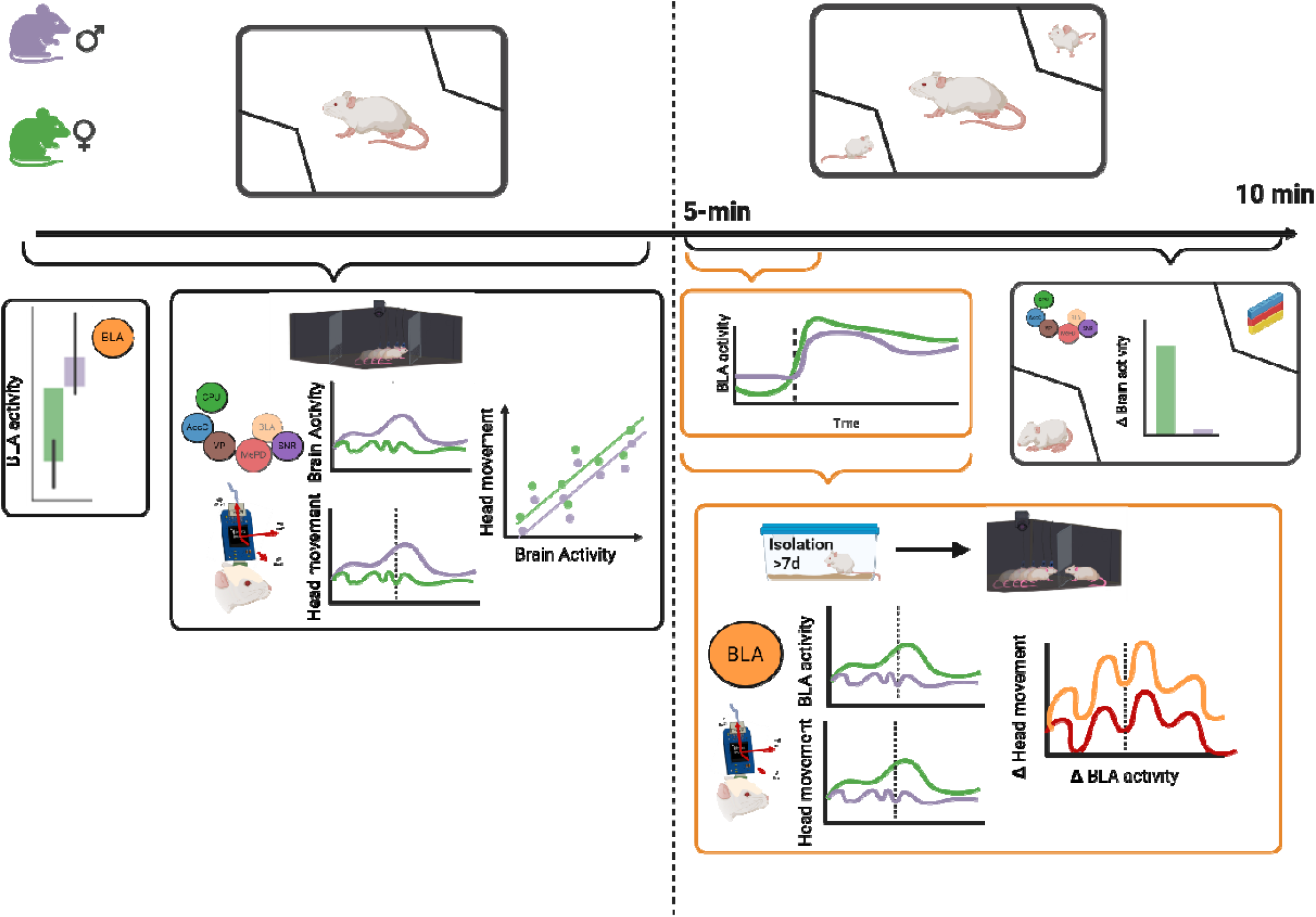

## Introduction

Social behaviour is essential for the survival of most mammalian species and is shaped by a combination of genetic, hormonal, and environmental factors. Sex represents a major biological variable influencing these processes, affecting how individuals perceive, evaluate, and respond to environmental cues [1–4]. Indeed, numerous studies in rodents have reported sex differences in behavioural strategies, including learning, decision-making, and exploration [5–13]. Social behaviour is no exception: sex differences have been observed across multiple paradigms, including social discrimination and preference tasks [14, 15]. In rats, only females exhibited a preference for interacting with a stressed cagemate over a naïve one [13]. Importantly, many neuropsychiatric disorders characterised by altered social behaviour show marked sex biases in prevalence. For example, autism spectrum disorder is more common in males, while anxiety and depression are more prevalent in females [19, 20].

Social decision-making involves multiple interacting processes, including the detection of social cues, evaluation of context, and integration of internal states such as motivation and arousal [15–19]. These processes unfold dynamically, requiring continuous updating as individuals navigate complex social environments. Consistent with this, the concept of the “social brain”, comprising a distributed network of interconnected brain regions, has emerged [20]. Understanding social behaviour, therefore, requires examining neural activity at the systems level. Notably, one section of the social brain which is well known to be involved in regulating social approach and avoidance is the amygdalo-striatal network, centered around the basolateral amygdala (BLA) and nucleus accumbens (AcbC) [21, 22].

Electrophysiological approaches using chronically implanted electrode arrays (EArs) enable simultaneous recordings from multiple brain regions in freely behaving animals, providing a powerful framework for studying large-scale neural dynamics [23]. Previous studies have shown that specific oscillatory patterns of local field potential (LFP) signals, particularly in the theta (4–12 Hz) and gamma (30–80 Hz) frequency bands, are modulated during social interactions and may reflect internal states such as attention and arousal [24–26]. Specifically, theta oscillations likely encode internal states like alertness, arousal and attention and may persist beyond social exposure, acting as a neural correlate of internal state [25]. However, most in vivo electrophysiological studies of social behaviour have focused on a single sex, typically males, or on individual brain regions, leaving sex-dependent, brain-wide dynamics largely unexplored.

Here, we address this gap by investigating sex differences in population neural activity across a distributed amygdalo-striatal network in freely behaving mice. Using chronically implanted EArs, we recorded LFP and multi-unit activity (MUA) simultaneously from multiple amygdalo-striatal regions during four distinct social discrimination tasks. By combining electrophysiological recordings with multimodal behavioural measurements, including video tracking and head acceleration, we examined how neural dynamics relate to behaviour in a sex- and context-dependent manner. This approach provides new insights into the network-level mechanisms underlying social behaviour and how they differ between males and females.

## Methods

### Animals

The subjects used in this study were adult (2–3 months old) male (N = 11) and female (N = 22) ICR (CD-1) mice, obtained from Envigo (Rehovot, Israel). Social stimuli included juvenile ICR mice (3–4 weeks old, used exclusively for the SP test) or adult male and female ICR mice of the same strain and age as the subjects. All animals were housed in the SPF mouse facility at the University of Haifa under controlled conditions (22± 2°C, 40-60% humidity) with a reverse 12-hour light–dark cycle (lights off at 7:00 a.m.). Food (standard chow; Envigo) and water were available *ad libitum*. All behavioural tests were conducted during the dark phase under dim red-light illumination. Stimulus mice were group-housed (3–5 per cage). Subject mice were housed individually for approximately one week following EAr implantation surgery to allow recovery and during the experiment to protect the implant. All procedures were approved by the Institutional Animal Care and Use Committee (IACUC) of the University of Haifa (protocols UoH-IL2203-139, 1077U).

### Electrode array

Following the protocol published in 2022 [23], we custom-designed electrode array (EAr) using 50 μm Formvar-insulated tungsten wires (50–150 kΩ, #CFW2032882; California Wire Company). The electrodes were arranged to target six brain regions: Nucleus Accumbens Core (AcbC; AP = +1.75 mm, ML = −1.25 mm, DV = −4.88 mm), Basolateral Amygdala (BLA; AP = −1.25 mm, ML = −3.25 mm, DV = −4.88 mm), Caudate Putamen (CPU; AP = −0.25 mm, ML = −3.25 mm, DV = −4.56 mm), Posterodorsal Medial Amygdala (MePD; AP = −2.25 mm, ML = −2.25 mm, DV = −4.88 mm), Substantia Nigra Reticulata (SNR; AP = −3.25 mm, ML = −1.25 mm, DV = −4.88 mm), and Ventral Pallidum (VP; AP = −0.25 mm, ML = −2.25 mm, DV = −4.88 mm). Before implantation, the EAr was coated with DiI (1,1’-Dioctadecyl-3,3,3’,3’-tetramethylindocarbocyanine perchlorate; #42364, Sigma-Aldrich) to allow for post-mortem verification of electrode tip placements. The brain areas recorded for each mouse are summarised in Additional file 1, Summary.

### Surgery

Subject mice were anaesthetised via intraperitoneal injection of a Ketamine–Domitor mixture (0.13 mg/g and 0.01 mg/g, respectively). Anaesthesia depth was regularly monitored using the toe-pinch reflex, and maintained with isoflurane (0.5–1%) delivered via a low-flow anaesthesia system (∼200 mL/min; SomnoFlo, Kent Scientific). Body temperature was kept constant at approximately 37 °C using a closed-loop, custom-built temperature controller. During the surgery, mice were placed in a stereotaxic frame (Kopf Instruments, Tujunga, CA) with the head in a flat position. The scalp was shaved and removed, and two anchor screws (0-80, 1/16”, M1.4) were implanted into the temporal bone. The craniotomy was performed by slow drilling into the skull. Coordinates for the target brain regions were marked over the left hemisphere, and the exposed brain surface was kept moist with sterile cold saline throughout the procedure. The reference and ground wires were inserted into designated burr holes at the back of the skull. The EAr was slowly lowered onto the exposed brain surface using a motorised micromanipulator (MP-200, Sutter Instruments). Once in place, the EAr and anchor screws were secured with dental cement (Unifast Trad). Post-surgery, animals received daily subcutaneous injections of Norocarp (0.005 mg/g) and Baytril (0.03 mL/10 g) for three days. They were given one week to recover before the start of the experimental procedures.

### Histological Validation

Subjects were transcardially perfused with 4% paraformaldehyde, and brains were post-fixed in 4% paraformaldehyde at 4 °C for 48 hours. Horizontal sections (50 µm thick) were cut using a vibratome (VT1200S, Leica). Electrode tracks were visualised by the fluorescent traces left by the DiI (red)-coated electrodes in DAPI-stained sections (Fluoroshield with DAPI, F6057, Sigma-Aldrich) using an epifluorescence microscope (Ti2 Eclipse, Nikon). Electrode placement was confirmed by aligning fluorescence images with the Mouse Brain Atlas [27].

### Behavioral Paradigms

Each mouse subject participated in four distinct social discrimination tests: Social Preference (SP), Sex Preference (SxP), Isolation-State Preference (ISP), and Stressed-State Preference (SSP), as previously described [12, 28–30]. The number of sessions and animals recorded and included in the analysis for each test are listed in Additional file 1. During each session, we recorded behaviour using a video camera, head-movement dynamics via an accelerometer (see below), and brain activity via the implanted EAr. Stimulus animals were always novel to the subject. In all tests except the SxP, stimuli were matched by sex. The SP test exposed the subject to a social stimulus (a novel, group-housed juvenile mouse) and an object (a Lego toy). In the SxP test, stimuli consisted of unfamiliar, group-housed adult male vs. female mice. In the ISP test, the subject encountered an isolated (for at least seven days) adult mouse versus a group-housed conspecific. For the SSP test, the subject encountered a stressed conspecific (restrained in a perforated 50-ml plastic tube for 15 minutes before the test) versus a naive, group-housed conspecific. Before each test session, subject mice were briefly exposed to isoflurane to minimise stress during the connection of the head-stage to the implanted EAr. Habituation was then conducted for 10 minutes, allowing the animal to acclimate to the arena containing empty stimulus chambers, before the onset of recording. Each recording session consisted of three consecutive 5-minute stages: a pre-test stage (with empty chambers placed at opposite corners of the arena), a test stage (with stimuli introduced into the chambers), and a post-test stage (with empty chambers again). Subject animals were tested twice daily over three consecutive days, with two sessions in the morning and two in the afternoon. The subject remained connected throughout the two sessions, with a 10-minute interval before each recording. Each subject completed three sessions of each of the four tests in a pseudo-randomised order (Fig. 1A). Sessions were excluded under the following conditions: if the head stage or connector detached from the subject, in cases of recording failure, or if the subject engaged in fewer than two bouts of stimulus investigation during the session (see Additional file 1). An investigation was defined as a continuous period lasting more than 2 seconds, starting when the subject touched the stimulus chamber and ending when the subject detached from the chamber. These exclusion criteria account for the variability in valid sessions and subjects across different tests (see Additional file 1).

**Figure 1.**
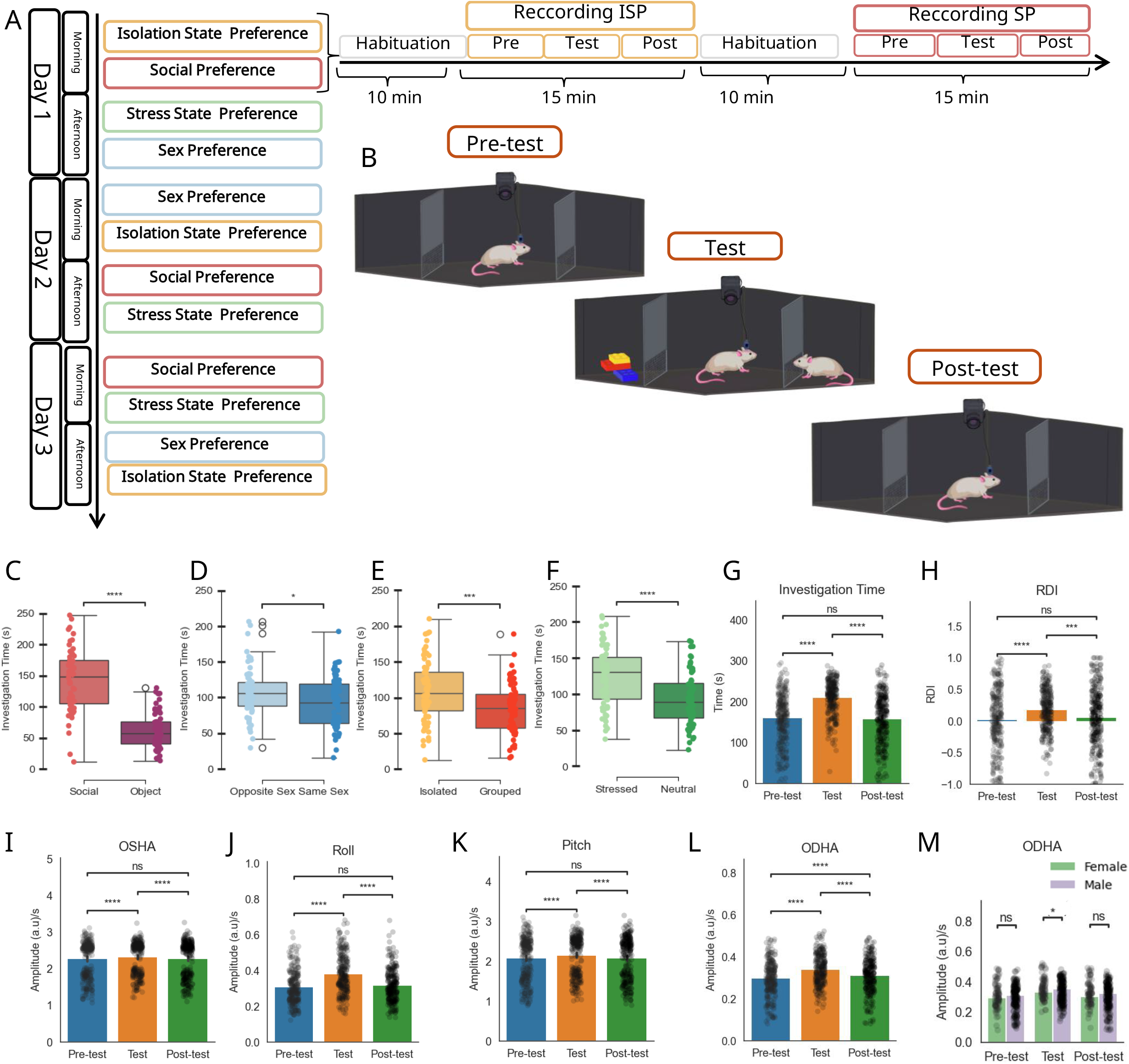
Behavioural features across the various stages of the social discrimination test. **A**. Timeline of the various tasks conducted by the recorded mice. **B**. Schematic representations of the setup at each stage (Pre-test, Test, Post-test). **C**. Mean investigation time was measured separately for each stimulus during the social preference (SP) test (t = 10.8, p < 0.001; t-test). **D**. As in (C) for the sex preference (SxP) test (t = 2.51, *p* = 0.013; t-test). **E**. As in (C) for the isolation-state preference (ISP) test (t = 3.78, *p* = 0.003; t-test). **F**. As in (C) for the stress-state preference (SSP) test (Females: t = 5.01, *p* < 0.001; t-test). **G**. Mean investigation time across stages (Pre-test, Test, and Post-test). Post hoc analysis following a main effect in a two-way mixed-model ANOVA for Stage: Pre-test vs Test (W = 284, p < 0.001), Test vs Post-test (W = 242, *p* < 0.001) (Wilcoxon test with FDR correction). **H**. Same as (G) for the RDI signal: Pre-test vs Test (W = 1285, p < 0.001), Test vs Post-test (W = 1334, *p* < 0.001) (Wilcoxon test with FDR correction). **I**. Same as (G) for the OSHA signal: Pre-test vs Test (W = 3504, *p* < 0.001), Test vs Post-test (W = 6521, *p* < 0.001) (Wilcoxon test with FDR correction). **J**. Same as (G) for the Roll signal: Pre-test vs Test (W = 4068, *p* < 0.001), Test vs Post-test (W = 4406, *p* < 0.001) (Wilcoxon test with FDR correction). **K**. Same as (G) for the Pitch signal: Pre-test vs Test (W = 9414, *p* < 0.001), Test vs Post-test (W = 9018, *p* < 0.001) (Wilcoxon test with FDR correction). **L**. Same as (G) for the ODHA signal: Pre-test vs Test (W = 3504, p < 0.001), Test vs Post-test (W = 6521, p < 0.001), and Pre-test vs Post-test (W = 11001, *p* < 0.001) (Wilcoxon test with FDR correction). **M**. Mean ODHA comparing Sex across stages. Post hoc analysis following a main effect in a two-way mixed-model ANOVA for Sex during test: (U = 8609, *p* = 0.039) (Mann-Whitney-Wilcoxon test with FDR correction).

### Behavioural Setups

The experimental setup has already been described [23, 28, 29]. It consisted of a black matte Plexiglas arena measuring 30 × 22 × 35 cm, which contained two triangular chambers (isosceles triangles with a 12 cm base and 35 cm height). The base of each triangular structure was covered with a metal mesh (18 cm × 6 cm, with 1 cm × 1 cm holes) to allow olfactory, visual, and limited tactile contact with stimulus animals.

These triangular stimulus chambers were placed in two randomly selected opposing corners of the arena. The entire arena was placed at the center of an acoustically isolated chamber that was electrically shielded and grounded to the electrophysiology recording system using 2-mm aluminium plates.

### Recordings and data analysis

#### Video

A high-resolution monochromatic camera (Flea3 USB3, FLIR—formerly Point Grey) was mounted at the top of the acoustically isolated chamber and connected to a computer. Behaviour was recorded at 30 frames per second using the FlyCapture2 software (FLIR). Subject behaviour was tracked using the TrackRodent algorithm for tethered mice (WhiteMouseWiredBodyBased3_7_19), as previously described [31]. Behavioural parameters, including investigation time, number and duration of investigation bouts and the Relative Discrimination Index (RDI), were calculated as previously described [28, 32, 33].

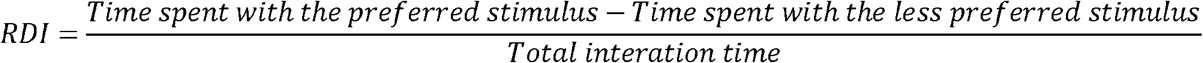

#### Head acceleration

Accelerometer recording and data analysis were conducted as previously described [34]. Briefly, head acceleration was recorded along three axes - surge, heave, and sway, corresponding to the X, Y, and Z axes, respectively, using an ADXL335 accelerometer (Analog Devices, Wilmington, MA), integrated into the Intan head-stages (see details below). Acceleration signals were sampled at 20 kHz and saved at 5 kHz. The accelerometer responds to both movement and the gravity vector. After calibrating the raw signal for each axis, we extracted four components. First, the calibrated signal was divided into dynamic acceleration by applying a band-pass filter (1-100 Hz) and static acceleration by applying a low-pass filter at 1 Hz. Then, the arcsine function was applied to the X and Z axes of the static acceleration data to calculate the inclination angles as follows:

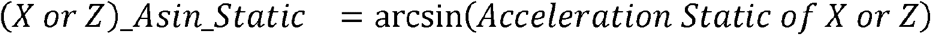

The arccos function was applied to the Y axis of the static acceleration data to calculate the inclination angle as follows:

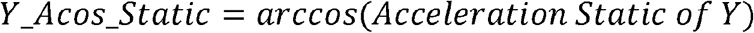

We then calculated the square root of the sum of squares for all three axes, which we termed OSHA and ODHA for the static and dynamic components, respectively.

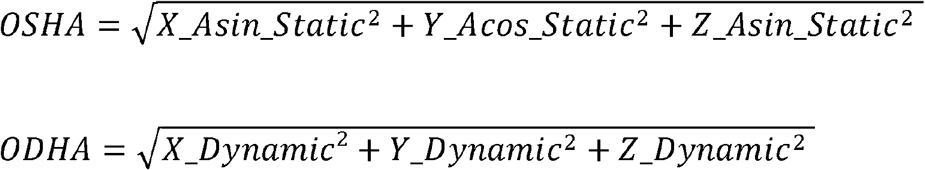

The Roll and the pitch were then derived from the Z and Y axes.

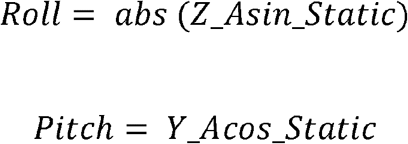

#### Electrophysiology

Subjects were connected to either 32- or 16-channel head-stages (Part #C3324 and Part #C3335, respectively; Intan Technologies, Los Angeles, CA) via a custom-made Omnetics-to-Mill-Max adaptor (Mill-Max model 852-10-100-10-001000). Electrophysiological signals were acquired using the RHD2000 evaluation system, with the head-stage connected through an ultra-thin SPI interface cable (Part #C3213; Intan Technologies). Recordings were sampled at 20 kHz. To ensure precise temporal alignment, we synchronised the video recordings via a TTL trigger pulse, and camera frame strobes were recorded using the system’s digital input channel.

Only brain regions recorded in more than five sessions across at least three mice were included in the analysis (Additional file 1). Electrophysiological signals were processed using custom-written code in MATLAB 2024a. The raw signals were downsampled to 5□kHz. LFP power was extracted at four frequency bands, including low frequencies (1–3□Hz), theta (4–12□Hz), gamma (30–80□Hz), and high frequencies (150–250□Hz) in two stages: first, we applied a low-pass filter at 300□Hz using a Butterworth filter. Second, we processed the signals according to the relevant time window. When calculating the LFP power along extended periods (e.g., 5 minutes), time–frequency estimates were computed with MATLAB’s *spectrogram* function with the following parameters: 2-second windows based on discrete prolate spheroidal sequences (DPSS), 50% overlap, frequency increments of 0.5□Hz, and time bins of 0.5□s. The final LFP power result was then averaged across the frequency band to express it in dB/Hz units. For calculating the LFP power along short epochs (e.g., several seconds), time–frequency decomposition was performed with an analytic Morlet (*amor*) wavelet transform [35] using the function *cwt* in MATLAB. We used the Morlet wavelet transform as it provides adaptive time–frequency resolution, allowing better characterisation of transient and non-stationary oscillatory events than the fixed-window Fourier transform [36]. The sampling frequency was 500 Hz, frequency range 1–250 Hz, and 15 voices per octave. Wavelet power was computed as |WT|^2^, and reported as relative power (dB/Hz).

Finally, to extract multi-unit activity (MUA), the raw electrophysiological signals were band-pass filtered between 0.3 kHz and 5 kHz using a Butterworth filter. Then, the time of spikes in each channel was determined using a threshold of 3.5 times the standard deviation calculated for each minute of the recording separately.

To avoid contamination from movement artefacts during stimulus insertion and removal, we excluded the 30 seconds preceding these events from all electrophysiological analyses.

From these calculated signals, we derived three types of measurements:

1. **Stage-based averages**: For each brain region and each 5-minute stage (pre-test, test, and post-test), we calculated the mean power for all 5 minutes within each frequency band and the mean MUA activity for each brain area. Changes were defined as the difference in mean power between the test and pre-test stages.
2. **First-minute analysis**: In some cases, we assessed changes in neural activity during the first minute following stimulus introduction. For that, signals were z-scored using the 60 seconds preceding stimulus insertion as the baseline. As explained above, we excluded the −30 to 0 second time window to avoid potential contamination from mechanical noise caused by the insertion of the stimuli.
3. **Event-aligned activity**: We also examined brain activity aligned to specific behavioural events - start and end of investigation bouts. At the beginning of the investigation, we z-scored the signal from −1 to +2 seconds relative to the moment the mouse touched the chamber, using the −2 to −1 second window as the baseline. Z-scoring was performed separately for each bout, and the resulting values were then averaged across all bouts corresponding to a given stimulus within each session. Importantly, only investigation bouts longer than 2 seconds were included to ensure meaningful engagement and reduce variability from short transient contacts.

#### PCA

We performed principal component analysis (PCA) using the five electrophysiological features mentioned above (four LFP power bands and MUA) across all recorded brain areas. The PCA was plotted separately for each brain region, first using data from the pre-test stage, and then using the changes in signals between the test and pre-test stages. This approach allowed us to visualise the data’s variance structure and examine how each brain region contributed to potential sex differences in neural activity.

### Decision tree classifier model

We trained multiple classification models using electrophysiological features to distinguish male and female mice. We used the mean values of five electrophysiological features in the first model during the pre-test stage. A second model was trained using the changes in mean power across the four frequency bands and MUA during the 5-minute test stage. Both models employed the *RandomForestClassifier* from the Scikit-learn package in Python, with 100 decision trees. We randomly downsampled the larger class when necessary to ensure balanced class representation. We used a cross-validation approach, leaving out at least 4 mice (and 16 sessions) for testing in each fold. Classifier performance was assessed via confusion matrices computed on the test sets. Feature importance was quantified as the mean decrease in Gini impurity, reflecting the extent to which each feature reduced classification error across decision splits. To evaluate whether the observed distributions were independent of sex, we applied a chi-square test of independence (*chi2_contingency*, SciPy).

### Statistical Analysis

All the data analyses were done using Python 3.11.11. All results and details of the statistical tests are summarised in Additional file 1.

Figures were created with the Seaborn package (versions 0.12.0 and 0.11.0), and statistical tests were performed using the Pingouin package (version 0.5.4) [37]. Normality was assessed using the Shapiro–Wilk test, homoscedasticity with Levene’s test, and sphericity with Mauchly’s test. Parametric tests were applied when these assumptions were met, including paired and independent t-tests, depending on the group dependence, and a two-way ANOVA for multi-group comparisons. For repeated-measures designs, either a one-way repeated-measures ANOVA or a mixed-model ANOVA was used, depending on the data structure. When assumptions for parametric testing were violated, non-parametric alternatives, such as the Wilcoxon signed-rank test, Mann–Whitney U test, Kruskal–Wallis test, and Friedman test, were used. We used a logistic regression model and Barnard’s exact test for binary data to assess whether the data distribution deviated from a random distribution. *Post hoc* analyses were performed following significant main effects, with multiple comparisons corrected using the False Discovery Rate (Benjamini & Hochberg method) via the Statannotations package. Associations between variables were assessed using Pearson’s correlation, and significance was evaluated via permutation testing using 10,000 shuffles. Sample sizes are summarised in Additional file 1.

## Results

### Multi-modal analysis reveals sex-specific murine social behaviour

Male and female ICR mice were recorded during the four behavioural tests described below, conducted over three days (two sessions in the morning and two sessions in the afternoon) in a pseudo-randomised order, as presented in Fig. 1A. In each session, the subject mouse was placed in an experimental arena containing two empty stimulus chambers located in randomly chosen opposite corners of the arena. After a 10-minute habituation stage, the recording session started with a 5-min pre-test stage, followed by a 5-minute test stage after replacement of the empty chambers with stimulus-containing chambers. At the end of the test stage, the stimulus-containing chambers were replaced by empty chambers, and a 5-minute post-test stage was recorded (Fig. 1B). Video clips from the various stages were analysed using TrackRodent [31], to extract, among other variables, the time dedicated by the subject to investigate each stimulus. These were used to calculate the Relative Discrimination Index (RDI), reflecting the preference of the subject to investigate one stimulus over the other [12, 23, 28]. As previously reported by us [12, 30], in the social preference (SP) test, subjects spent more time (preferred) investigating a novel sex-matched juvenile conspecific (social stimulus) than a Lego toy (object stimulus) (Fig. 1C; t = 10.8, p < 0.001; t-test). In the sex preference (SxP) test, subject mice preferred an age-matched conspecific of the opposite sex over a same-sex social stimulus (Fig. 1D; t = 2.51, *p* = 0.013; t-test independent). In the isolation-state preference (ISP) test, subjects preferred an isolated (7-14 days) mouse over a group-housed mouse (Fig. 1E; t = 3.783, *p* < 0.001; t-test independent). Finally, in the stress state preference (SSP) test, subject mice preferred a stressed mouse over the non-stressed (naive) mouse (Fig. 1F; t = 5.01, *p* < 0.001; t-test independent). Notably, none of these tests showed a main effect of sex (Fig. S1A, D, G, J).

Next, we examined behavioural changes across the various stages of the session, regardless of the test type. For that, we used the total time the animal spent investigating each of the two stimuli and the RDI. We also included in this analysis four fine-grained head movement features extracted from an accelerometer device present on the electrophysiological recording headstage, as previously described by us [34]: Overall Dynamic Head Acceleration (ODHA), Overall Static Head Acceleration (OSHA), Roll and Pitch.

Notably, for each of the six variables mentioned above, values were higher during the test stage than during both the pre-test and post-test stages (Fig. 1G–L; Investigation time: F = 162.7, *p* < 0.001; RDI: F = 11.7, *p* < 0.001; OSHA: F = 12.9, *p* < 0.01; Roll: F = 91, *p* < 0.001; Pitch: F = 22.3, *p* < 0.001; ODHA: F = 107.5, *p* < 0.001; two-way mixed-model), suggesting that these features genuinely reflect the animal’s social behavior. However, only one out of these six variables, ODHA, showed a main effect of sex (Fig. 1M and Fig. S2A-E; ODHA: F = 4.11, p = 0.046, two-way mixed-model ANOVA), with a higher level of ODHA in males specifically during the test stage (Fig. 1M; Test: U = 8609, *p* = 0.039, Mann-Whitney-Wilcoxon test with FDR correction). Since the ODHA reflects the animal’s rapid head movement [34], these sex differences in ODHA, which appear specifically during the test stage, may suggest that males’ social investigation behaviour is more energetic than that of females.

### Network-level electrophysiological dynamics in a subset of amygdalo-striatal regions are modulated by social context and sex

To explore sex differences in the electrophysiological dynamics across the social brain at the system level, we used custom-built EArs (Fig. 2A) to record Local Field Potentials (LFP) and multi-unit activity (MUA) simultaneously from a subset of amygdalo-striatal regions during the behavioural experiments described above. Due to some variability in targeting accuracy, not all regions were recorded in every subject (electrode-tip placements were verified *post mortem*; Fig. 2B). These brain regions included the Nucleus Accumbens Core (AcbC), Caudal Putamen (CPU), Ventral Pallidum (VP), Basolateral Amygdala (BLA), Posterodorsal Medial Amygdala (MePD), and the Substantia Nigra Reticulata (SNR) (Fig. 2C).

**Figure 2.**
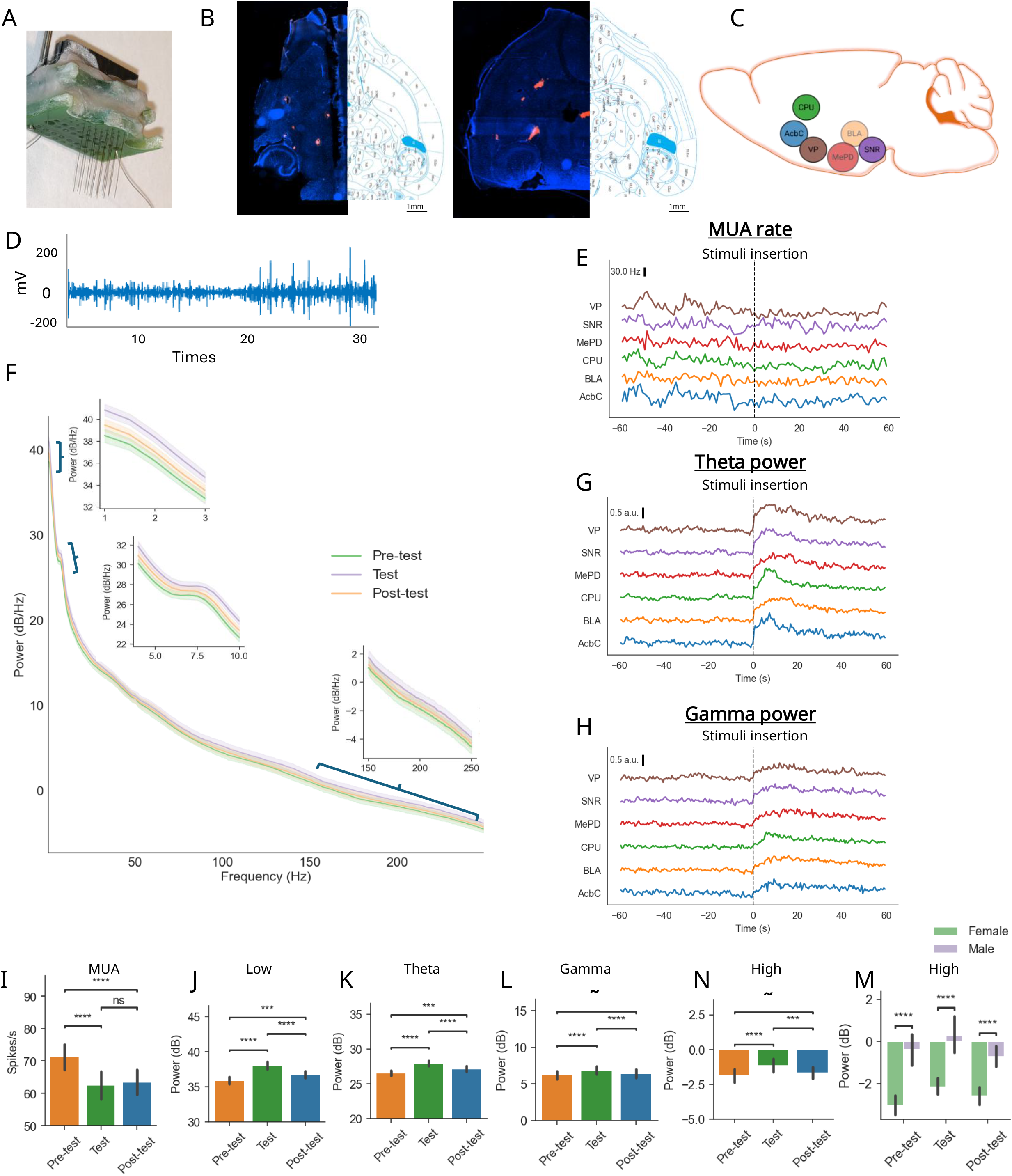
Electrophysiological features analysed across stages and sex. **A**. Image of an electrode array. **B**. Example of horizontal brain slice images at −3.60 mm (left) and −4.28 mm (right) depth from bregma, stained with DAPI. The electrode track is labelled with DiI. **C**. Map of the targeted brain areas: AcbC – Accumbens core, BLA – Basolateral amygdala, CPU – Caudate putamen, MePD – Posterodorsal Medial amygdala, SNR – Substantia nigra reticulata. **D**. Raw trace illustrating spike detection in the CPU of a female mouse. **E**. Mean MUA traces from all brain areas, showing the last minute of the Pre-test (before the vertical line) and the first minute of the Test for each brain area. **F**. Average (± SEM) PSD profile of the CPU of female mice: Pre-test (green), Test (purple), and Post-test (orange), with a zoom on the low-, theta-, and high-frequency ranges. **G**. Same as (E), but filter for the LFP theta band. **H**. Same as (E), but filter for the LFP gamma band. **I**. Mean (± SEM) MUA across stages for all brain regions and all four tests. Post hoc analysis following a main effect in a two-way mixed-model ANOVA for Stage: Pre-test vs Test (t = 9.25, p < 0.001), Test vs Post-test (t = 8.43, *p* < 0.001) (paired t-test with FDR correction). **J**. Mean (± SEM) power of the low frequency band from all brain regions across the stages in all four tests. Post hoc analysis following a main effect in a two-way mixed-model ANOVA for Stage: Pre-test vs Test (t = –13.4, *p* < 0.001), Test vs Post-test (t = 10, *p* < 0.001), and Pre-test vs Post-test (t = –4, *p* < 0.001) (paired t-test with FDR correction). **K**. Same as (J), but for the theta band (4–10 Hz). Stage: Pre-test vs Test (W = 21, *p* < 0.001), Test vs Post-test (W = 157, *p* < 0.001), and Pre-test vs Post-test (W = 26, *p* = 0.006) (Wilcoxon test with FDR correction). **L**. Same as (J), but for the gamma band (30–150 Hz). Pre-test vs Test (W = 156, p < 0.001), Test vs Post-test (W = 215, *p* < 0.001), and Pre-test vs Post-test (t = 402, *p* = 0.057) (Wilcoxon test with FDR correction). **N**. Same as (J), but for the high-frequency band (150–250 Hz). Pre-test vs Test (W = 139, *p* < 0.001), Test vs Post-test (W = 236, *p* = 0.002), and Pre-test vs Post-test (t = 427, *p* = 0.01) (Wilcoxon test with FDR correction). **M**. Mean (± SEM) power of the high frequency band across Stage and Sex for all brain regions for all four tests. Post hoc analysis following a main effect in a two-way mixed-model ANOVA for Sex: Pre-test (U = 76, *p* < 0.001), Test (U = 77, *p* < 0.001), Post-test (U = 75, *p* < 0.001) (Mann-Whitney-Wilcoxon test with FDR correction).

Figure 2D shows raw traces of MUA recorded from the CPU in a female mouse. In general, a decrease in MUA was observed following stimulus introduction. As apparent in Fig. 2E, which shows the mean traces of MUA rate for each recorded brain region, averaged across all mice, during the last minute of the pre-test and the first minute of the test. Following stimulus introduction, a reduction in firing rate was observed consistently across all brain regions.

Figure 2F shows the power spectrum density (PSD) calculated for LFP signals recorded in the CPU of females, as an example. We chose to investigate four frequency bands: theta (4–10 Hz) and gamma (30–150 Hz), which have been frequently studied in the context of social behaviour [33, 38], as well as the less-explored low (1–3 Hz) and high (150–250 Hz) frequency bands. Unlike MUA, mean traces of LFP theta (Fig. 2G) and gamma (Fig. 2H, S3) power revealed an apparent increase in power at the first minute of the test, across all brain regions.

When we examined overall changes between the three stages averaged across all six brain areas, we observed a main effect of stage for all five electrophysiological features. Specifically, MUA rate decreased during both the test and post-test stages, compared to the pre-test stage (Fig. 2I; MUA: F = 11.6, *p* < 0.001; one-way repeated-measures ANOVA). In contrast, LFP power across all frequencies was higher during the test stage compared to both the pre-test and post-test stages (Fig. 2J–N; LFP– low: F = 99.3, *p* < 0.001; LFP–theta: F = 37.99, *p* < 0.01; LFP-gamma: F = 11.10, *p* = 0.002; LFP–high: F = 11.23, *p* = 0.002; one-way repeated-measures ANOVA). These results show that the social behaviour induced a general change in electrophysiology across the entire brain-region network we recorded from. In addition, the high-frequency band (150–250 Hz) showed a main effect of sex (Fig. 2M; F = 6.7, p = 0.049; two-way mixed-model ANOVA), with males consistently showing higher power than females across all stages.

Together, these findings demonstrate that social interaction (social context) modulates neuronal spiking and LFP oscillatory power across multiple regions in the amygdalo-striatal section of the social brain, with frequency-specific sex effects.

### Sex-specific electrophysiological differences are observed already before the test

The sex-specific differences in the power of high-frequency LFP oscillations during the pre-test stage (Fig. 2M) suggest that sex-specific electrophysiological patterns exist already before the test stage. To further examine this possibility, we performed a principal component analysis (PCA) on the data recorded during the pre-test stage using the five electrophysiological features described above and plotted the data separately for each brain region (Fig. 3A–F). In some areas, such as the AcbC, CPU, MePD, and VP, we observed a clear separation between male and female mice, suggesting that the variance in these regions is strongly influenced by sex already before the test. As expected, we observed no apparent separation of the data due to the test that will follow the pre-test stage, in any of the brain regions (Fig. S4A-F).

**Figure 3.**
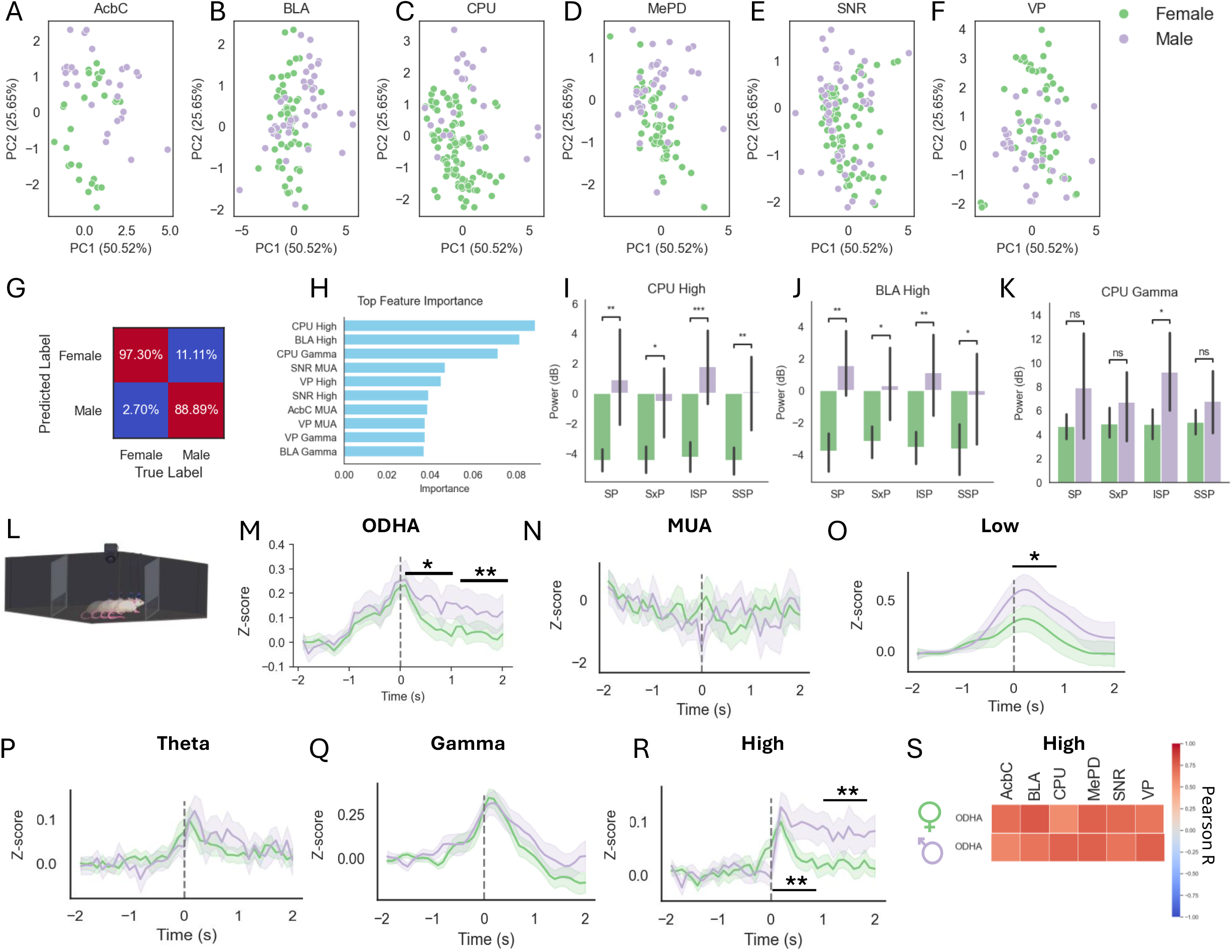
Brain activity and behavioural sex differences during the pre-test stage. **A–F**. PCA was performed on the electrophysiological features of all brain areas (see methods). The first two principal components (PC1 and PC2) explained 47.48% and 27.7% of the total variance, respectively. Each point represents an individual session, colour-coded by sex (female in green, male in purple). Each brain area was plotted separately: (A) accumbens core (AcbC), (B) basolateral amygdala (BLA), (C) caudate putamen (CPU), (D) posterodorsal medial amygdala (MePD), (E) substantia nigra reticulata (SNR), and ventral pallidum (VP). **G**. Confusion matrix for a binary Random Forest classifier model predicting sex from electrophysiology-derived data across all tests during the 5 min of the pre-test stage. Each category’s percentage is displayed at the center of each cell (Female: *χ* = 16.09, *p <* 0.001; Male: χ = 13.63, *p* <0.001; Chi-square test). **H**. Bar plot showing the top 10 features ranked by importance in the predictive model. Features are ordered from highest to lowest importance. **I**. Mean (± SEM) power of the CPU high-frequency band (150 to 250 Hz) across test types and sex. Post hoc analysis following a main effect in a two-way ANOVA for sex: SP (U = 13, *p* = 0.003), SxP (U = 20, *p* = 0.016), ISP (t = 17, *p* < 0.001), and SSP (t = 19, *p* = 0.003) (Mann-Whitney-Wilcoxon test with FDR correction). **J**. Same as (I) but for the BLA high-frequency band (150 to 250 Hz): SP (t = –4.3, *p* < 0.001), SxP(t = –4.2, *p* = 0.007), ISP (t = –3.67, *p* = 0.001), and SSP (t = –4.26, *p* = 0.042) (t-test with FDR correction). **K**. Same as (I) but for gamma power in CPU (30 to 150 Hz): ISP (U = 38, *p* = 0.011) (Mann-Whitney-Wilcoxon test with FDR correction). **L**. Schematic representation of a mouse approaching an empty chamber. **M**. Mean (± SEM) z-scored ODHA signal two seconds before and after the start of investigation bouts across all sessions for females (green) and males (purple). Time 0 represents the start of the bout; the baseline was defined as the interval from −2 to −1 s. Post hoc Wilcoxon tests with FDR correction were performed following a main effect in a two-way mixed-model ANOVA for sex (0 to 1 s: W = –2.33, *p* = 0.021; 1 to 2 s: W = –2.60, *p* = 0.01). **N**. Mean (± SEM) z-score of MUA from all brain areas as in (M). **O**. Mean (± SEM) z-score of low-frequency LFP signal from all brain areas as in (M). Post hoc Wilcoxon tests with FDR correction following a main effect in two-way mixed-model ANOVA (0 to 1 s: W = 49, *p* = 0.028). **P**. Mean (± SEM) z-score of theta-frequency LFP signal as in (M). **Q**. Mean (± SEM) z-score of gamma-frequency LFP signal as in (M). **R**. Mean (± SEM) z-score of high-frequency LFP signal as in (M). Post hoc Wilcoxon tests with FDR correction following a main effect in two-way mixed-model ANOVA (0 to 1 s: W = 61, *p* = 0.0096; 1 to 2 s: W = 51, *p* = 0.003). **S**. Correlation matrix between ODHA at bout start (−1s to 2s) and mean high-frequency LFP power in each brain area. The correlation scale is shown to the right.

To quantify this effect, we implemented a random forest classifier to predict the animal’s sex, based on the electrophysiological features. This classifier performed significantly better than chance, correctly identifying ∼89% of the female data and 75% of the male data (Fig. 3G; Females: χ^2^ = 16.09, *p* < 0.001; Males: χ^2^ = 13.63, *p* < 0.001). We then extracted the most important features driving this classification and identified three top-ranked features that showed significant sex differences (Fig. 3H). *Post hoc* analyses revealed that male mice exhibited significantly higher high-frequency power in both the CPU (F = 74.9, *p* < 0.001; two-way ANOVA) and BLA (F = 38.4, *p* < 0.001; two-way ANOVA) across all tests. They also showed higher gamma power in the CPU (F = 14.9, *p* < 0.001; two-way ANOVA), although this was only significant in the ISP test, with a trend observed in all other tests (Fig. 3I-K). Together, these results suggest that during the pre-test stage, before the social interaction, male and female mice already exhibit differences in specific electrophysiological features, particularly in selected brain regions. These differences likely reflect inherent sex-specific network states that may shape subsequent responses to social stimuli. Since mice underwent the test multiple times, we cannot exclude the possibility that the anticipation of the upcoming social interaction influenced their brain activity during the pre-test stage.

None of the behavioural features presented in Figure 1 showed significant differences between males and females before the test stage. However, in a previous publication [34], we reported sex differences in head acceleration when mice approached an empty chamber during the pre-test stage (see schematic depiction in Fig. 3L), with males exhibiting a greater increase in ODHA than females (Fig. 3M; F = 7.6, *p* = 0.01, two-way ANOVA). Based on this, we applied a similar analysis to the electrophysiological data by averaging the signal across the six recorded brain regions for each feature and aligning it with the stimulus investigation behaviour.

Signals were z-scored across the −2 to +2 seconds relative to the contact with the chamber, using −2 to −1 second as a baseline. We found that male mice displayed a significantly greater increase in both low-frequency and high-frequency power when touching an empty chamber (Fig. 3O, R; Low: F = 9.1, *p* = 0.03; High: F = 21.3, *p* < 0.001; two-way mixed-model ANOVA). In contrast, no significant sex differences were observed in theta or gamma bands (Fig. 3P, Q) or for MUA rate (Fig. 3N). For high-frequency power, the higher level observed in males as the subject started touching the chamber persisted for the first two seconds of touching the chamber, similar to the ODHA signal. Additionally, we observed a significant correlation between ODHA and high-frequency power across all brain areas in both sexes (Fig. 3S), whereas no such correlation was found for low-frequency power (Fig. S4G). Since we previously suggested [34] that ODHA could serve as an indicator of arousal state in mice, the high-frequency activity may similarly reflect arousal.

Altogether, these findings reveal that male and female mice exhibit distinct electrophysiological signatures even before social interaction begins, particularly in specific brain regions and frequency bands. Specifically, when approaching an empty chamber, male mice exhibited more rapid head movements and a higher level of high-frequency LFP oscillations compared to females. These sex differences, evident during the pre-test stage, suggest the existence of sex-specific basal neural states across the brain that may influence the following social interaction.

### Sex differences in brain electrophysiology during social interaction are distributed across features and brain areas

To examine sex differences related to social behaviour during the test stage, we computed the change (Δ) from baseline in each of the five electrophysiological features by subtracting the mean value calculated for the 5-minute pre-test stage from the one calculated for the 5-minute test stage. We first examined the general social context, irrespective of the test type, by performing a PCA, as described above for the pre-test stage. Unlike the pre-test stage, we did not observe a clear sex-dependent separation for any of the brain areas during the test stage (Fig. S5A–F). Nevertheless, when applying a random forest classifier using the same variables as for the pre-test stage, the model was able to classify sex, correctly identifying ∼86% of the female data and ∼72% of the male data (Fig. S5G; Female: χ^2^ = 5.13, *p* = 0.021; Male: χ^2^ = 4.63, *p* = 0.03). Yet, when applying two-way ANOVA to each brain area and electrophysiological feature individually, none reached statistical significance (Additional file 1: Figure S5), which explains why the importance scores were very similar across features. Thus, although the random forest model could successfully classify sex, indicating undeniable differences in the data, these differences seem to be distributed across brain areas and electrophysiological features, unlike in the pre-test stage.

Then, we averaged the electrophysiological features across all brain areas and looked for sex differences in a test-specific manner. With the exception of MUA, we found a significant interaction between sex and test type for all features (Additional file 1; Figure 4; LFP –low: F = 4.64, *p* = 0.02; LFP – theta: F = 5.47, *p* < 0.01; LFP –gamma Hz: F = 12.59, *p* < 0.001; LFP – high: F = 11.71, *p* < 0.001; two-way repeated-measures ANOVA). For MUA, we observed a trend toward significance (Additional file 1, F = 3.26, *p* = 0.051; two-way repeated-measures ANOVA).

**Figure 4.**
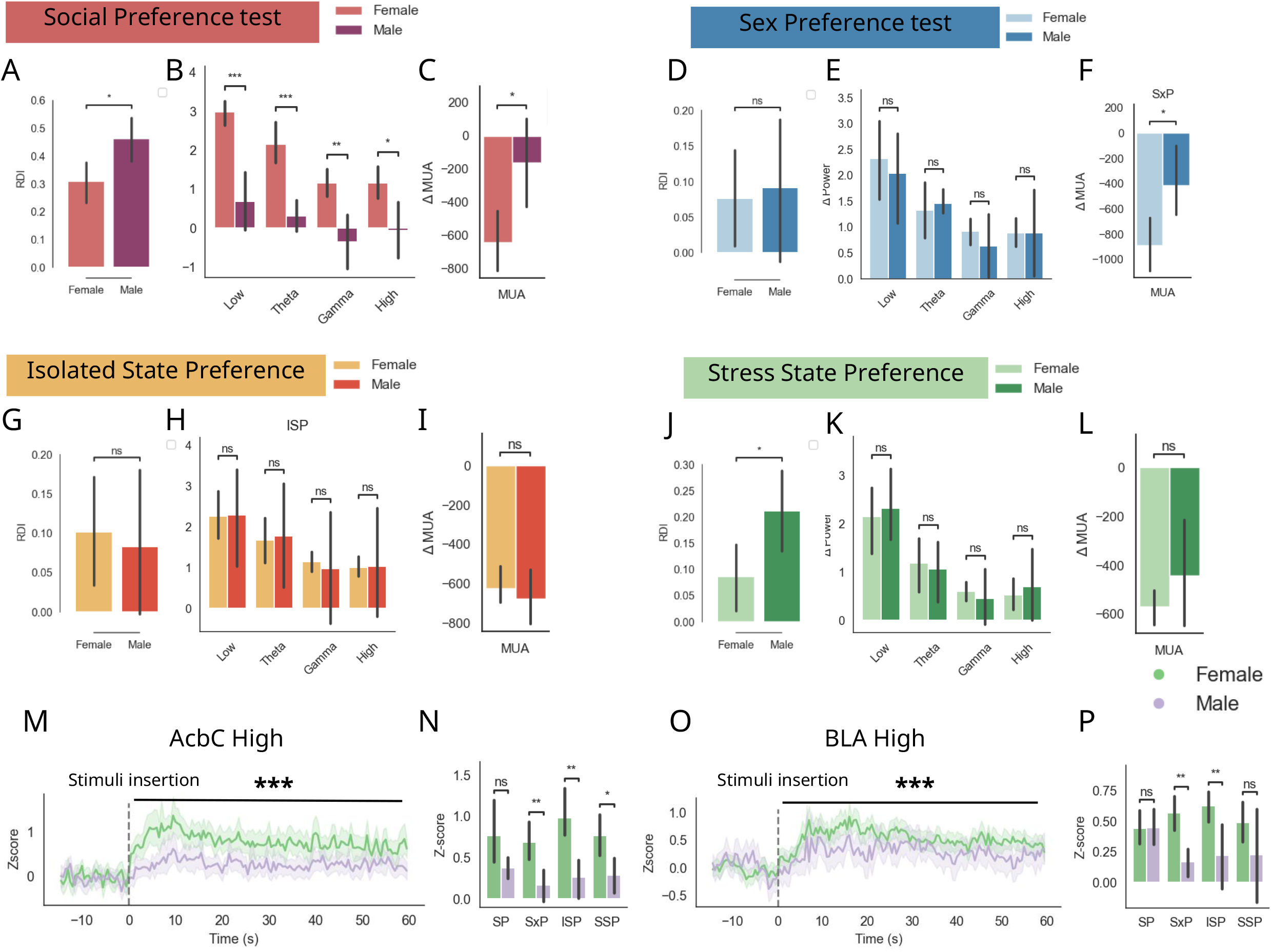
Context- and sex-dependent changes in brain activity during the various social discrimination tests. **A**. Mean (± SEM) RDI for each sex during the SP test (t = 2.54, *p* = 0.015; t-test). **B**. Mean (± SEM) LFP Δ power across four frequency bands (low, theta, gamma, high) between sexes during the SP test across all brain regions. Post hoc analysis following a main effect of Sex in a two-way mixed-model ANOVA: low frequency band (t = 4.97, *p* < 0.001), theta band (t = 5.66, *p* < 0.001), gamma band (t = 4.81, *p* < 0.001), high frequency band (t = 3.80, *p* = 0.003) (t-tests with FDR correction). **C**. Mean (± SEM) Δ MUA across all brain regions during the SP test for the two sexes (t = 2.59, *p* = 0.030; t-test). **D**. Same as (A), for the SxP test. **E**. Same as (B), for the SxP test. **F**. Same as (C), for the SxP test (t = 2.50, *p* = 0.033; t-test). **G**. Same as (A), for the ISP test. **H**. Same as (B), for the ISP test. **I**. Same as (C), for the ISP test. **J**. Same as (A), for the SSP test (t = –2.24, *p* = 0.025; t-test). **K**. Same as (B), for the SSP test. **L**. Same as (C), for the SSP test. **M**. Mean (± SEM) z-score of AcbC high-frequency power (150–250 Hz) across all brain areas, from 15 seconds before to 60 seconds after stimulus insertion, for each test. Curves represent the average across all sessions for females (green) and males (purple). Time 0 marks the insertion of stimuli; baseline was defined as the 1-minute interval preceding stimulus insertion. A two-way ANOVA with FDR correction revealed a main effect of Sex (F = 11.92, *p* < 0.001). **N**. Mean (± SEM) AcbC high-frequency power across Stage and Sex for all brain areas. Post hoc analysis following a main effect in a two-way mixed-model ANOVA showed significant differences in: SxP (t = 3.13, *p* = 0.01), ISP (t = 3.4, *p* = 0.005), and SSP (t = 4.0, *p* = 0.031) (independent t-tests with FDR correction). **O**. Same as (M), for BLA high-frequency power. A main effect of Sex was found (F = 31.7, *p* < 0.001). **P**. Same as (N), for BLA high-frequency power. Significant effects in SxP (U = 39, *p* = 0.008) and ISP (U = 48, *p* < 0.008) (Mann-Whitney-Wilcoxon test with FDR correction).

Surprisingly, although males showed a stronger behavioural preference for the social stimulus in the SP test than females (Fig. 4A; t = 2.54, *p* = 0.011; independent t-tests), the mean change in LFP power was significantly higher in females (Fig. 4B; F = 76.71, p < 0.001; two-way repeated-measures ANOVA). *Post hoc* analyses revealed that this sex difference was significant across all four frequency bands (low, theta, gamma, and high). Similarly, MUA activity also changed more in females than males, with a significantly stronger decrease observed in females (Fig. 4C; T = 2.59, p = 0.032; paired t-test). Moreover, when we examined whether there was any change in brain activity across the 5-minute test stage in males, none of the electrophysiological features differed significantly from zero (Fig. 4B–C).

For the SxP and ISP tests, where the RDI was similar between sexes (Fig. 4D, G), we found no difference in the electrophysiology change (Fig. 4E, F, H, I), except for a stronger decrease in MUA rate during the test stage for females compared to males (Fig. 4F; t = 2.5, *p* = 0.033; independent t-tests).

Finally, for the SSP test, male mice showed a stronger preference toward the stressed mouse than females (Fig. 4J; SSP: t = 2.25, *p* = 0.025; independent t-tests). However, all electrophysiology features were similar between male and female (Fig. 4K, L). Thus, when looking for changes in brain electrophysiology across the 5-minute test stage, we did not find any clear correlation between such changes and the animal’s social behaviour. However, this approach does not account for the dynamic and robust changes in neural activity that occur at the beginning of social interaction, immediately following stimulus insertion (see Fig. 2D, F, G). To examine this, we z-scored the LFP power and MUA rate using the last minute of the pre-test stage as a baseline and analysed the changes occurring during the first minute of the test stage, separately for each electrophysiological feature and brain area.

Two features showed a significant effect of sex (when tested for the effects of Sex and Test): high-frequency power in the AcbC and BLA (Fig. 4M, O; AcbC: F = 31.26, *p* < 0.001; BLA: F = 11.98, *p* < 0.001; two-way ANOVA with FDR correction). In the AcbC, high-frequency power increased significantly more in females than males during the SxP, ISP, and SSP tests following the insertion of stimuli (Fig. 4N), and a similar trend was observed in the SP test. In the BLA, females exhibited a significantly higher increase in high-frequency power during the first minute of the SxP and ISP tests, but not in the SP and SSP tests (Fig. 4P). Notably, no significant sex differences were observed when examining the means of the various kinematic variables across the first minute of each test (Fig. S6A–D).

Altogether, these findings suggest that sex-specific electrophysiological differences in the first minute of the test stage, the most intensive period of the social interaction, are accompanied primarily by high-frequency power in the AcbC and BLA, especially during the SxP and ISP tests.

### BLA activity as a neural correlate for sex-dependent early social approach behaviour

We then examined how preference dynamics evolved over time across sexes, for each test. ISP was the only test that revealed a clear Sex × Time interaction (Fig. 5A and Fig. S7A-C; F = 3.92, p = 0.004; two-way mixed-model ANOVA). During the first minute of the test, males preferred the isolated stimulus, whereas females did not. This difference was consistent with the greater change in high-frequency BLA power observed in females during the first minute following stimulus insertion (Fig. 4O). These results led us to explore the relationship between the electrophysiological and behavioural variables during the ISP at a higher time resolution. Specifically, we examined the electrophysiological dynamics during approach behaviour to either the isolated or the group-housed stimulus. For three frequency bands (theta, gamma, and high-frequency), LFP power in the BLA increased more in females than in males, specifically when the animals approached the isolated stimulus (BLA theta: F = 5.75, p = 0.023; BLA gamma: F = 5.56, p = 0.026; BLA high: F = 9.13, p = 0.005; two-way mixed-model ANOVA). This sex effect was observed only in the BLA (Additional file 1: “1^st^ min – each BrainArea”), and was almost undetectable when animals approached the group-housed stimulus, with a significant sex main effect observed only in the high-frequency band and only in the ANOVA test (F = 5.27, p = 0.03), with no significant *post hoc* differences (Fig. 5B–D). The sex-specific increase in BLA activity (particularly theta and gamma) was not observed during the last 4 minutes of the test (Fig. S8A–B), thus paralleling the RDI sex differences (Fig. 5A). Only for the high-frequency band, the sex-dependent difference persisted during the later minutes (Fig. S8C; F = 4.90, p = 0.035), although the effect was weaker.

**Figure 5.**
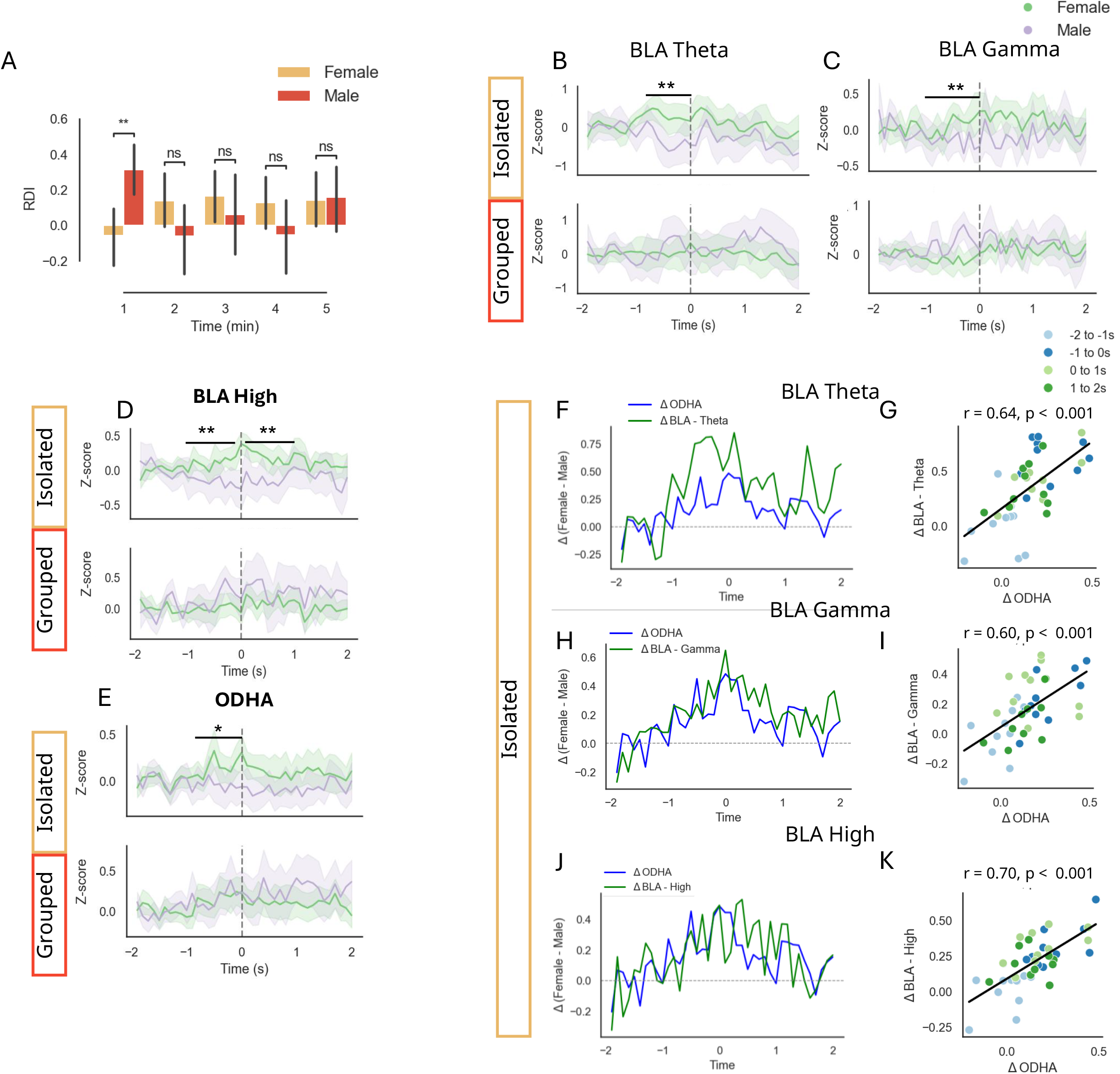
Behavioural sex differences correlate with BLA electrophysiological activity during early social exposure. **A**. Mean RDI (± SEM) across sex and time during the ISP test. Post hoc analysis following a main effect in a two-way mixed-model ANOVA revealed significant differences for Time: First minute (U = 358, *p* = 0.005) (Mann-Whitney-Wilcoxon test with FDR correction). **B**. Mean (± SEM) z-score of BLA theta power from 2 seconds before to 2 seconds after the onset of investigation of the isolated (top) and grouped (bottom) stimuli, for females (green) and males (purple), during the first minute of the ISP test. Time 0 marks the start of the bout; the baseline was defined from −2 to −1 s. Post hoc t-tests for independent samples with FDR correction revealed significant sex differences for the isolated stimulus from −1 to 0 s (t = 2.94, *p* = 0.007). **C**. Same as (B), for BLA gamma power. Isolated stimulus: −1 to 0 s (t = 2.78, *p* = 0.01), 0 to 1 s (t = 2.6, *p* = 0.015) (t-tests with FDR correction). **D**. Same as (B), for BLA high-frequency power (150–250 Hz). Isolated stimulus: −1 to 0 s (t = 3.07, *p* = 0.005), 0 to 1 s (t = 3.1, *p* = 0.005) (t-tests with FDR correction). **E**. Same as (B), but for ODHA signal. Isolated stimulus: −1 to 0 s (t = 2.1, *p* = 0.043) (t-tests with FDR correction). **F**. Time traces of ΔODHA (z-score Female ODHA minus z-score Male ODHA, in blue) and ΔBLA theta power (z-score Female minus Male, in green) aligned to the onset of approach to the isolated stimulus. **G**. Scatter plot showing the relationship between ΔODHA (x-axis) and ΔBLA theta power (y-axis) during investigation bouts with the isolated stimulus. Each point represents a time point, with colour indicating time within the event. Regression lines show the fitted linear model. The correlation was assessed using Pearson’s correlation coefficient, and significance was evaluated via permutation testing using 10,000 shuffles. Pearson correlation: r = 0.64, *p* < 0.001. **H**. Same as (F) but for gamma power. **I**. Same as (G), but for gamma power. Pearson correlation: r = 0.60, *p* < 0.001. **J**. Same as (F) but for the high-frequency power. **K**. Same as (G), but for high-frequency power. Pearson correlation: r = 0.70, *p* < 0.001.

When analysing head kinematics during approaches toward the stimuli, we found a significant sex effect for ODHA, with females showing greater head movement during the first minute (Fig. 5E). Similar to the LFP power, this difference was not present during the last four minutes (Fig. S8D). Notably, no sex differences in ODHA were observed when mice approached the group-housed stimulus. Moreover, during the first minute, when the subject began interacting with the isolated stimulus, the sex-difference in ODHA strongly correlated with the sex difference in BLA power across the theta, gamma, and high-frequency bands (Fig. 5F–K theta: r = 0.64, p < 0.001; gamma: r = 0.60, p < 0.01; BLA high: r = 0.70; Pearson correlation). However, during interaction with the grouped stimulus, these correlations dropped markedly (Fig. S9F; high: r = 0.42, p = 0.008; Pearson correlation and permutation testing), or disappeared (Fig. S9A-B, D; theta: r = 0.38, p = 0.014; gamma: r = –0.20, p = 0.223; Pearson correlation).

Altogether, these findings reveal a significant correlation between sex differences in behaviour and neural dynamics during the ISP test. Specifically, the elevated BLA power in females was most pronounced when approaching the isolated stimulus and strongly correlated with the ODHA, suggesting sex-dependent coordination between BLA and head movement during social exploration. Importantly, the sex differences in both behaviour and electrophysiology were limited to the beginning of the ISP test, thus showing social context dependency.

## Discussion

This study provides an overview of sex-specific behavioural and electrophysiological differences observed in ICR mice across multiple social discrimination tests. Using a multi-modal approach, we captured a rich behavioural profile and revealed that sex differences in social behaviour are both context-dependent and dynamic. These results are consistent with previous studies from our lab [12, 30] that demonstrated sex differences in the social behaviour of mice. Notably, the only test where we observed dynamic sex differences in the RDI was the ISP test; unlike males, females did not show a preference for the isolated conspecific during the first minute, while such a preference appeared later. In a previous study, we showed that C57BL/6 mice tested in the same paradigm also show sex differences, with no preference exhibited by females, in contrast to males [30]. Thus, the sex-dependent behavioural differences in the ISP test seem genuine and robust and could be explained either by a difficulty in recognising an isolated stimulus mouse or by a different response to it, exhibited by subject females.

### Pre-test sex-specific patterns of network activity may reflect sex-dependent internal state

Our recordings revealed sex differences in brain activity across several social discrimination tests. While we did not find sex-dependent behavioural differences during the pre-test stage, our findings indicate that sex-specific neural dynamics were present already before social interactions. This was particularly evident in the power of high-frequency LFP oscillations recorded in the BLA and CPU, as well as gamma-band LFP power in the CPU (Fig. 3I–K). Although mice may have anticipated the social encounter that follows the pre-test stage due to repeated sessions conducted with the same subject mouse, we found that the type (social context) of the following test did not influence the electrophysiological signals. These pre-test electrophysiological differences may reflect internal states induced in the subject by its placement in the experimental arena, such as anxiety or alertness. Alternatively, they may reflect differences in anticipation of a forthcoming social interaction.

Evidence from patch-clamp studies supports the existence of intrinsic sex differences in excitability across various brain regions. For instance, in the MPOA, female neurons are more excitable, show higher input resistance and exhibit lower firing threshold, whereas male neurons are larger and display stronger rebound activity [39]. In the anterior cingulate cortex (ACC), male mice exhibit stronger synaptic long-term depression than females, despite similar long-term potentiation across sexes [40]. Male mouse neurons fire at higher rates in the locus coeruleus, while female neurons show distinct passive membrane properties and proteomic profiles linked to protein synthesis and stress signalling [41]. Altogether, these studies suggest that neurons have intrinsic sex differences in their activity that depend on the brain area. Such differences may contribute to the sex differences in network electrophysiological activity observed by us during the pre-test stage.

### High-frequency LFP oscillations and sex differences

Theta and gamma LFP oscillations in the brain were most often studied in the context of social behaviour. During social encounters, most limbic brain regions exhibit higher theta and gamma power compared to baseline [24, 25]. Moreover, several mouse models of autism spectrum disorder (ASD) exhibit higher LFP theta and gamma rhythmicity than wild-type mice, even at rest [24, 29]. In our study, however, the sex differences were most prominent at high frequencies (150-250 Hz). This frequency band was the only band showing sex differences across all stages, when all tests were pooled together (Fig. 2M). It also exhibited sex differences at the beginning of stimulus investigation, depending on the specific context and brain area (Fig. 3R, Fig. 4O-P and Fig. 5D). High-frequency oscillations (≈ 200 Hz) occur not only in the hippocampus during sleep and memory processes but also across cortical and subcortical regions in rodents, including the entorhinal cortex, mPFC, neocortex, and amygdala [42–45]. Cortical ripples are associated with large neuronal depolarisations [46, 47], boosting firing rates [43]. Similarly, amygdalar High-frequency oscillations entrain the firing of many neurons [45], with a higher proportion showing phase-locking than rate modulation [48]. Moreover, these oscillations are highly coherent across distant brain sites [45]. High-frequency oscillations, therefore, represent a possible mechanism for explaining the organisation of social networks at the brain-wide level and may account for a large part of the emerging sex differences.

Similar to the electrophysiological features, while multiple behavioural features showed sex differences at specific time points, the ODHA showed such differences most consistently. Moreover, the ODHA sex differences tended to correlate with multiple electrophysiological features and most consistently with high-frequency LFP power. For example, both ODHA and high-frequency LFP power were higher in males when averaged across the entire test stage (Fig. 1M and Fig. 2M). These findings reinforce our previously published hypothesis that ODHA may serve as a proxy for internal states [34]. Interestingly, ODHA sex differences became specifically associated with the BLA signals at distinct time windows, mainly during the first minute of social interaction.

### A pivotal role of the BLA in sex differences during social interactions

Unlike the relatively focused sex differences in network activity observed during the pre-test stage, those observed during the full 5-minute test period were distributed across the network, rather than being dominated by 1-2 brain areas. At specific time points, certain brain regions and frequency bands emerged as primary contributors to these differences. One area that stood out, showing consistent sex differences across time points, was the BLA.

During the first minute of the SxP and ISP tests, females showed stronger increases in BLA high-frequency LFP power following stimulus presentation. Moreover, BLA LFP power across several frequency bands was higher, specifically in females, when interacting with the isolated stimulus during the first minute of the ISP session. This dynamic and flexible pattern of sex-dependent BLA activity could not be explained by pre-existing sex differences, as it was male BLA high-frequency power that was higher during the 5-minute pre-test. Together, these results point to the BLA as a key contributor to sex-dependent neural activity during social interaction. Importantly, in the ISP test, female ODHA was higher than male ODHA, specifically during the first minute of interaction with the isolated stimulus, mirroring the pattern of sex differences in BLA activity.

Thus, BLA activity exhibits strong context- and time-dependence. The dynamic sex-dependent preference observed in females during the ISP test was accompanied by a more substantial increase in BLA high-frequency power during the first minute of the test. This difference between males and females was highly correlated over time with the sex-dependent difference in ODHA. Notably, this pattern was observed specifically when females initiated an interaction with the isolated, but not with the grouped stimulus. One may speculate that females are more cautious at the beginning of interaction, showing higher BLA activity and less interaction with the isolated stimulus animal. In contrast, this behaviour is more stable in males, where we also do not see the fluctuation in BLA activity.

The BLA plays a central role in regulating social interactions in rodents by integrating emotionally salient sensory information with behavioural responses [49]. Experimental studies have shown that the BLA contributes to social recognition, social preference, and the evaluation of the emotional states of conspecifics [50].

This brain area also exhibits sexual dimorphism, as the density of dopaminergic synaptic boutons in murine BLA, as well as the density of cells, is significantly higher in the male brain than in the female brain [51, 52]. In accordance, we found the BLA to be a key player in the dynamic sex differences observed in the amygdalo-striatal network during social interactions.

### Limitations

Although we observed correlations between neural sex differences and behaviour, particularly during the approach toward an isolated stimulus, our study remains inherently correlational. Future work using causal approaches, such as chemogenetic and optogenetic manipulations, will be necessary to determine whether the observed neural differences directly contribute to the behavioural patterns reported here.

In addition, precisely targeting the same brain areas across different animals and across sexes remains technically challenging. Consequently, our dataset did not allow for systematic comparisons of functional connectivity between regions through measures such as coherence. Investigating potential sex differences in inter-regional connectivity, therefore, represents an important direction for future research.

Furthermore, the electrophysiological recordings analysed here do not allow us to distinguish between specific neuronal cell types. Future studies combining large-scale recordings with cell-type–specific approaches will be important to determine which neuronal populations contribute to the observed effects.

Finally, the behavioural paradigm used in this study relied on relatively controlled conditions. Extending these investigations to more complex and naturalistic paradigms involving groups of freely moving animals may provide further insights into how these neural dynamics relate to behaviour in ecologically relevant contexts.

### Perspectives and Significance

Our findings provide new insights into the neural correlates of sex differences in social behaviour and reveal that sex shapes both baseline and interaction-driven neural activity across the social brain network. Specifically, it suggests a pivotal role for high-frequency oscillations in the BLA for such sex differences. Future studies could investigate how manipulating these oscillations may influence sex differences in social behaviour, hence establishing a causal role between them.

## Conclusions

BLA neural activity, especially high-frequency oscillations, exhibited pronounced sex-specific, context- and time-dependent dynamics. These neural differences were associated with sex differences in social behavior and movement dynamics, particularly during the initial phase of social interaction. Together, our findings highlight the BLA as a key node underlying sex-specific dynamics of social behaviour.

## Supplementary Information

### Additional files

Additional file 1

- File format: .xlsx
- Title of data: Supplementary tables
- Description of data: Excel file containing supplementary tables associated with the analyses presented in this study.

Additional file 2

- File format: .docx
- Title of data: Supplementary figures
- Description of data: Word document containing the supplementary figure associated with this study.

## Acronyms

AcbC: Accumbens Core
ASD: Autism Spectrum Disorder
BLA: Basolateral Amygdala
Ear: Electrode array
HFOs: High Frequency Oscillations
ISP: Isolated State Preference
MePD: Posterodorsal Medial Amygdala
ODHA: Overall Dynamic Head Acceleration
OSHA: Overall Static Head Acceleration
PCA: Principal Component Analysis
PSD: Power Spectrum Density
SNR: Substantia Nigra Reticulata
SP: Social Preference
SSP: Stress State Preference
SxP: Sex Preference
RDI: Relative Discrimination IndexVP: Ventral Pallidium

## Declarations

### Ethics approval and consent to participate

All experimental procedures were approved by the Institutional Animal Care and Use Committee of the University of Haifa (UoH-IL2203-139, 1077U).

### Consent of publication

All authors read and approved the final manuscript.

### Availability of data and materials

The datasets supporting the conclusions of this article are available in the Zenodo repository: female data at https://zenodo.org/records/19565987 [53] and male data and analysis code at https://zenodo.org/records/19682930 [54].

### Authors’ contributions

A.P.: Formal analysis, Investigation, Methodology, Validation, Visualisation, Writing— original draft, and Writing— review & editing; S.N.: Data curation, Project administration, Software, Validation, Visualisation, Writing—original draft, and Writing—review & editing. S.W.: Conceptualisation, Funding acquisition, Project administration, Resources, Supervision, Writing—original draft, and Writing—review & editing.

## Acknowledgements

Some parts of the figures have been created using BioRender.com.

