## Additional file 2 for "Sex differences in neural activity across amygdalo-striatal network during social behaviour"

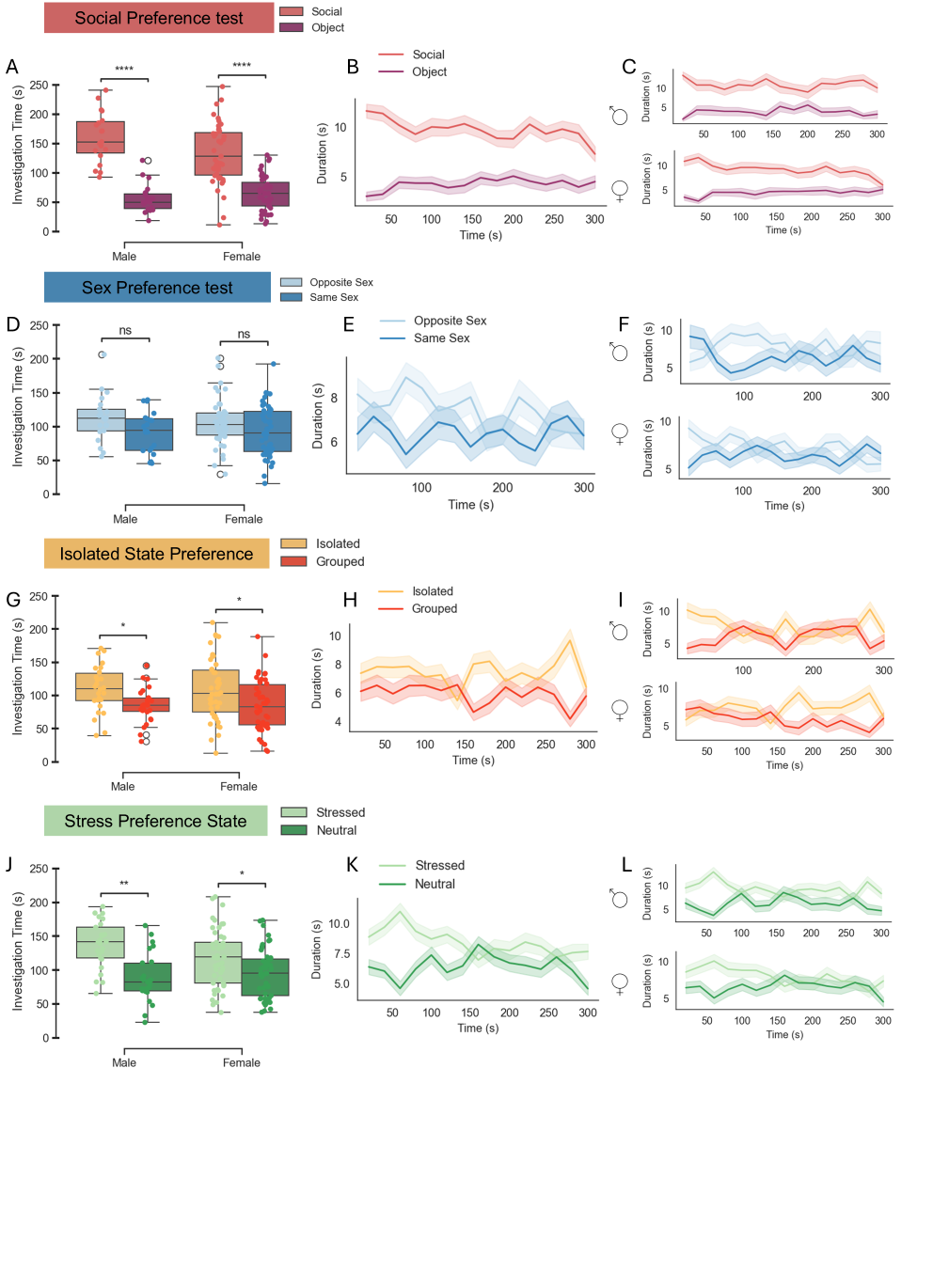

**Figure S1.**

**A.** Mean (± SEM) investigation time was measured separately for each stimulus during the social preference (SP) test for females (t = 6.48, p < 0.001) and males (t = 7.15, *p* < 0.001) mice.
**B.** Time trace of the duration spent with each stimulus (in seconds) over the 5-min SP test, pooled for both males and females together.
**C.** Same as (B), but separately for males (upper panel) and females (lower panel).
**D.** Same as (A) for the sex preference (SxP) test (Females: t = 1.39, *p* = 0.17; Males: t = 1.61, p = 0.122).
**E.** Same as (B) but for the SxP test.
**F.** Same as (E), but separately for males (upper panel) and females (lower panel).
**G.** As in (A) for the isolation-state preference (ISP) test (Females: t = 2.44, *p* = 0.019; Males: t = 2.45, p = 0.021).
**H.** Same as (B) but for the SxP test.
**I.** Same as (H), but separately for males (upper panel) and females (lower panel).
**J.** As in (A) for the stress-state preference (SSP) test (Females: t = 2.40, *p* = 0.024; Males: t = 3.68, p = 0.0018).
**K.** Same as (B) but for the SSP test.
**L.** Same as (K), but separately for males (upper panel) and females (lower panel)

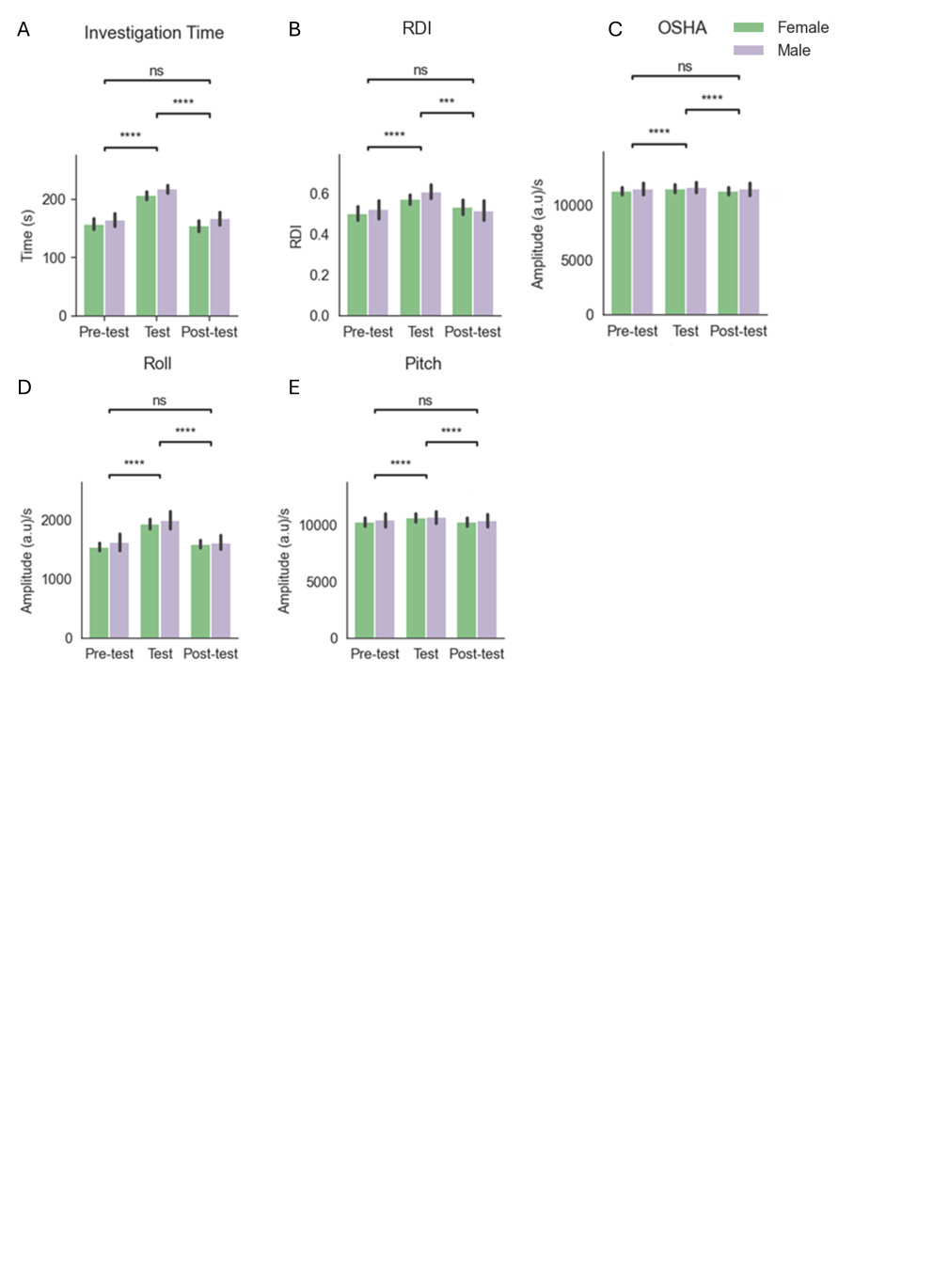

**Figure S2.**

**A.**  Mean investigation time across stages (Pre-test, Test, and Post-test). Post hoc analysis following a main effect in a two-way mixed-model ANOVA for Stage: Pre-test vs Test (W = 284, p < 0.001), Test vs Post-test (W = 242, *p* < 0.001) (Wilcoxon test with FDR correction).
**B.** Same as (A) for the RDI signal: Pre-test vs Test (W = 1285, p < 0.001), Test vs Post-test (W = 1334, *p* < 0.001) (Wilcoxon test with FDR correction).
**C.** Same as (A) for the OSHA signal: Pre-test vs Test (W = 3504, *p* < 0.001), Test vs Post-test (W = 6521, *p* < 0.001) (Wilcoxon test with FDR correction).
**D.** Same as (A) for the Roll signal: Pre-test vs Test (W = 4068, *p* < 0.001), Test vs Post-test (W = 4406, *p* < 0.001) (Wilcoxon test with FDR correction).
**E.** Same as (A) for the Pitch signal:: Pre-test vs Test (W = 9414, *p* < 0.001), Test vs Post-test (W = 9018, *p* < 0.001) (Wilcoxon test with FDR correction).

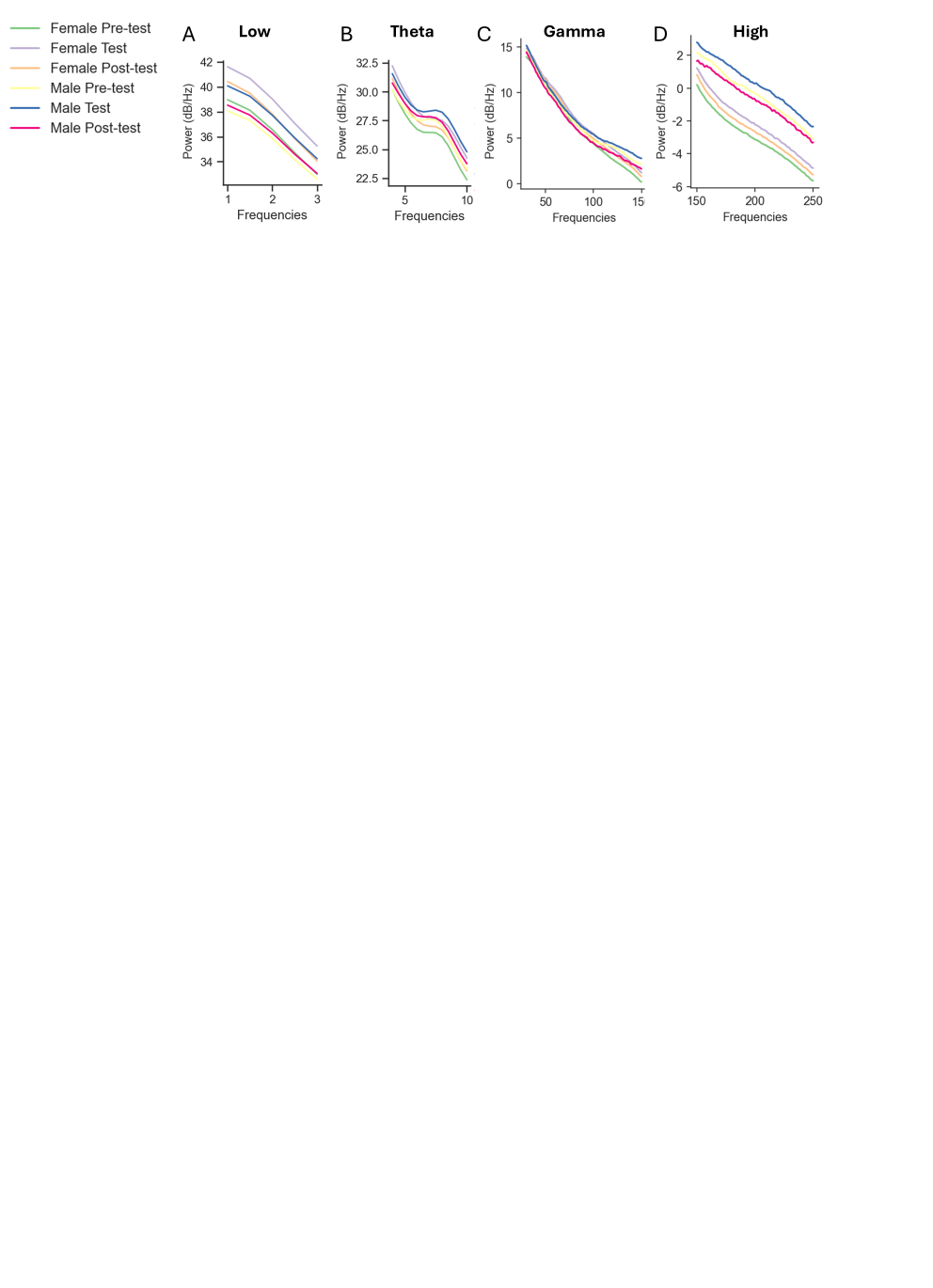

**Figure S3.**

**A.** Average PSD profile from low frequency LFP band: female Pre-test (green), female Test (purple), female Post-test (orange), male Pre-test (yellow), male Test (blue), male Post-test (red).
**B.** Same as (A), but for the LFP theta band.
**C.** Same as (A), but for the LFP gamma band.
**D.** Same as (A), but for the LFP high-frequency band.

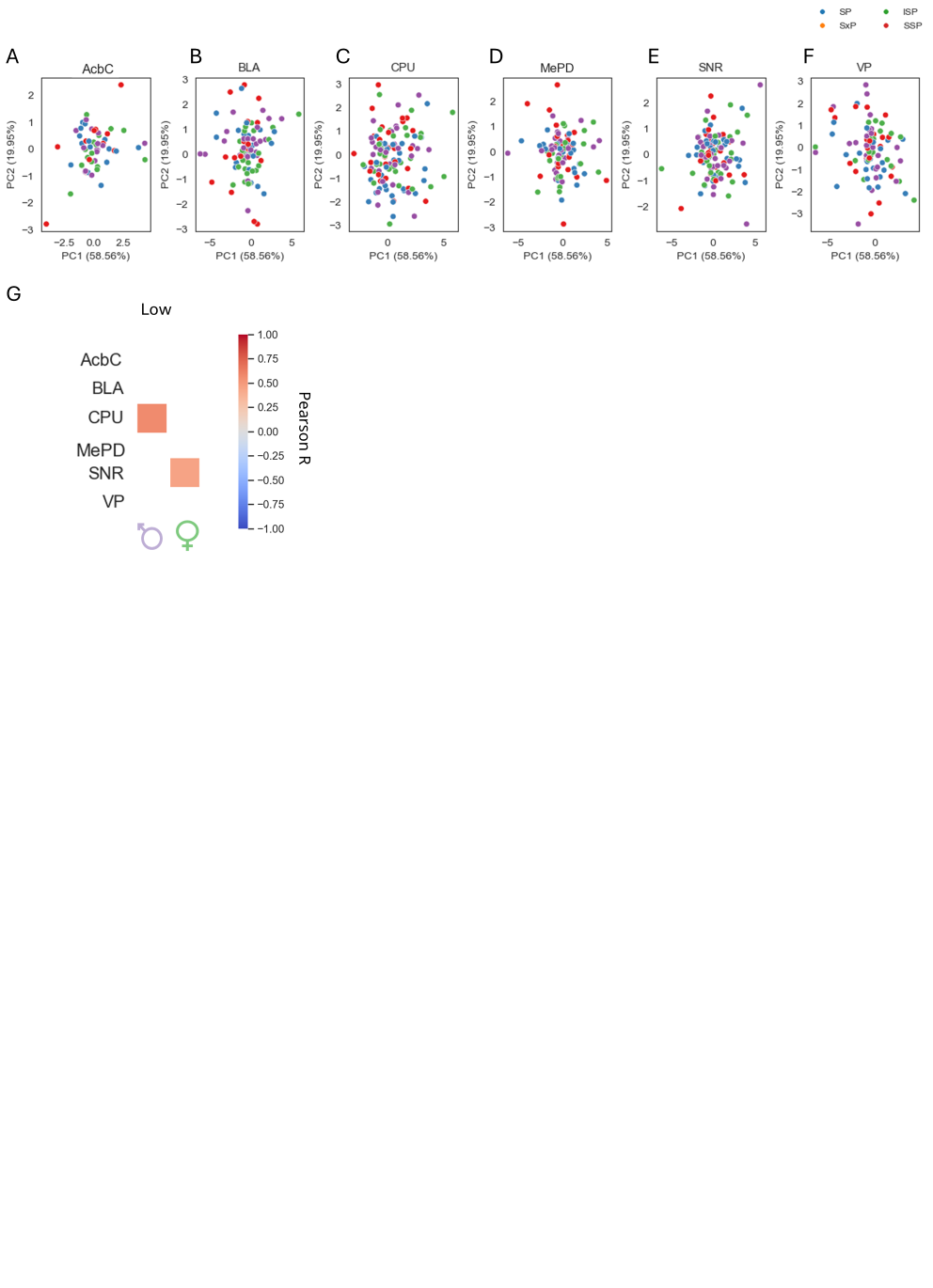

**Figure S4.**

**A–F.** PCA was performed on electrophysiological features of all brain areas. The first two principal components (PC1 and PC2) explained 47.48% and 27.7% of the total variance, respectively. Each point represents an individual session, colour-coded by test (SP in blue, SxP in orange, ISP in green, and SSP in red). Each brain area was plotted separately: (A) Accumbens core (AcbC), (B) basolateral amygdala (BLA), (C) caudate putamen (CPU), (D) posterior-dorsal medial amygdala (MePD), (E) substantia nigra reticulata (SNR), and (F) ventral pallidum (VP).
**G.** Correlation matrix between ODHA at bout start (-1s to 2s) and mean low LFP power (1 to 3 Hz) in each brain area. The correlation scale is shown on the right.

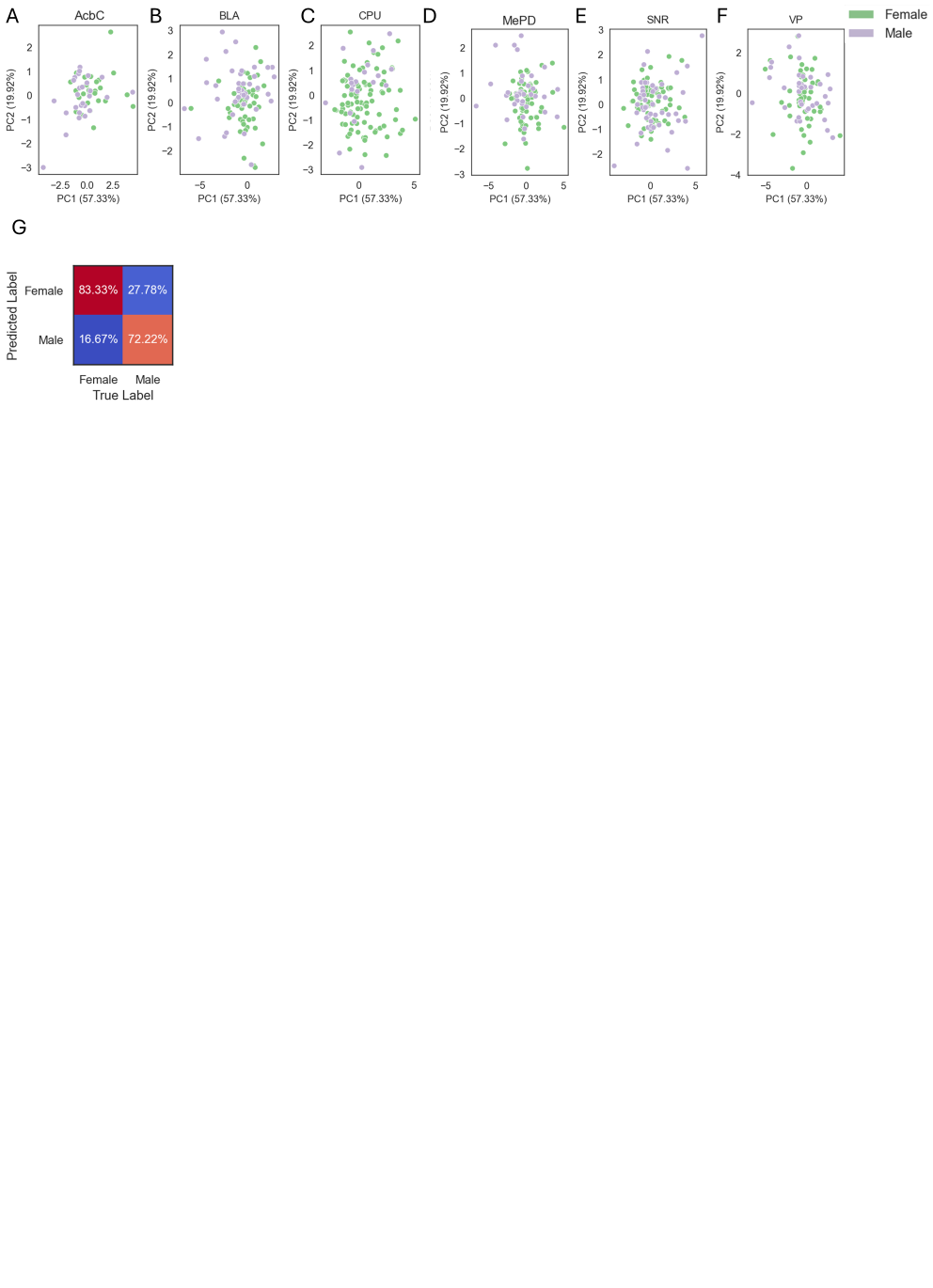

**Figure S5.**

**A–F.** PCA was performed on the Δ electrophysiology features of each brain area (mean electrophysiology feature during test stage minus mean electrophysiology during the pre-test stage. The first two principal components (PC1 and PC2) explained 57.33% and 19.92% of the total variance, respectively. Each point represents an individual session, colour-coded by sex (female in green, male in purple). Each brain area was plotted separately: (A) Accumbens core (AcbC), (B) basolateral amygdala (BLA), (C) caudate putamen (CPU), (D) posterior-dorsal medial amygdala (MePD), (E) substantia nigra reticulata (SNR), and (F) ventral pallidum (VP).
**G.** Confusion matrix for a binary Random Forest classifier model predicting sex from Δ electrophysiology-derived data across all tests during the 5 min of the pre-test stage. Each category's percentage is displayed at the center of each cell (Female: *χ* = 5.3, *p =* 0.021; Male: χ = 4.63, *p =* 0.03; Chi-square test).

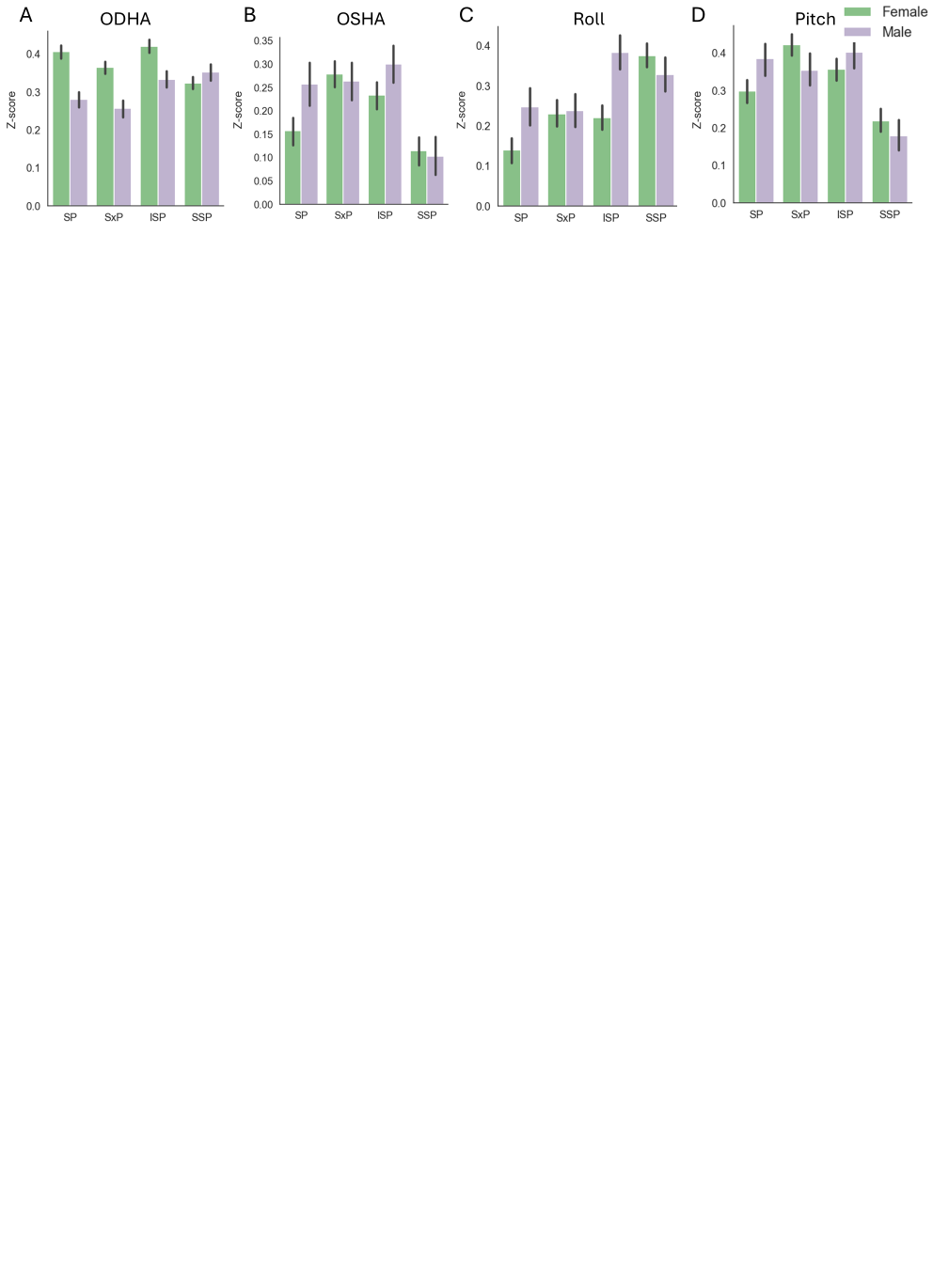

**Figure S6.**

**A.** Mean (± SEM) Z-scored ODHA for the first minute of interaction across Stage and Sex. Baseline was defined as the 1-minute interval preceding stimuli insertion.
**B.** Same as (A) but for the OSHA signal.
**C.** Same as (A) but for the roll signal.
**D.** Same as (A) but for the pitch signal.

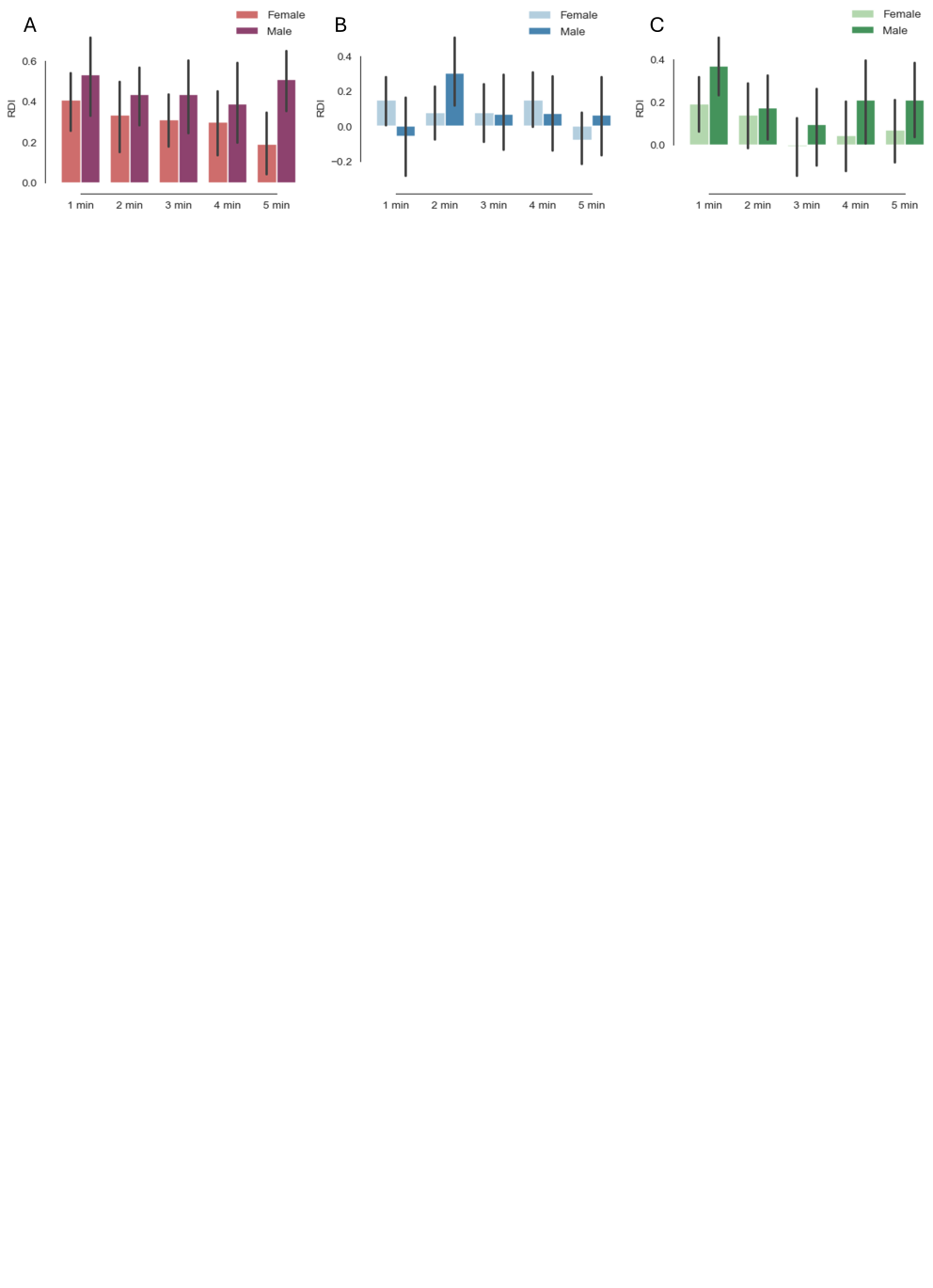

**Figure S7.**

**A.** Mean RDI (± SEM) across sex and time during the SP test.
**B.** Same as (A) but for the SxP test.
**C**. Same as (A) but for the SSP test.

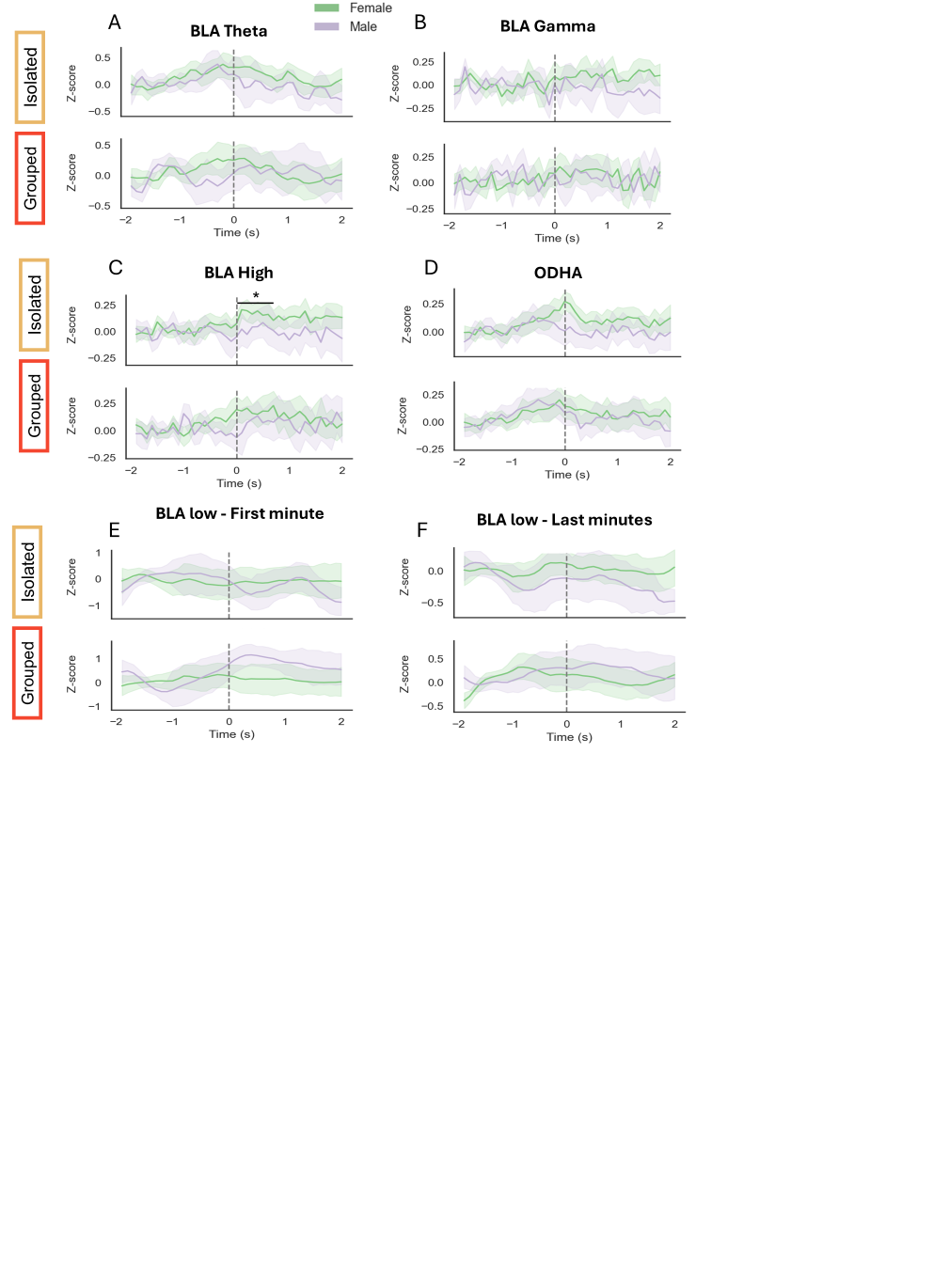

**Figure S8.**

**A.** Mean (± SEM) Z-score of BLA LFP theta power from 2 seconds before to 2 seconds after the onset of investigation of the isolated (top) and grouped (bottom) stimuli, for females (green) and males (purple), during the last four minutes of ISP test. Time 0 marks the start of the bout; the baseline was defined from -2 to -1 s.
**B.** Same as (A) for BLA LFP gamma power.
**C.** Same as (A) for BLA LFP high-frequency power. Isolated stimulus: 0 to 1 s (t = 2.10, *p* = 0.044) (t-tests with FDR correction).
**D.** Same as (A) for the ODHA signal.
**E.** Same as (A) but for the BLA LFP low-frequency power during the first minute of the test.
**F.** Same as (A) but for BLA LFP low-frequency power during the last four minutes of the test.

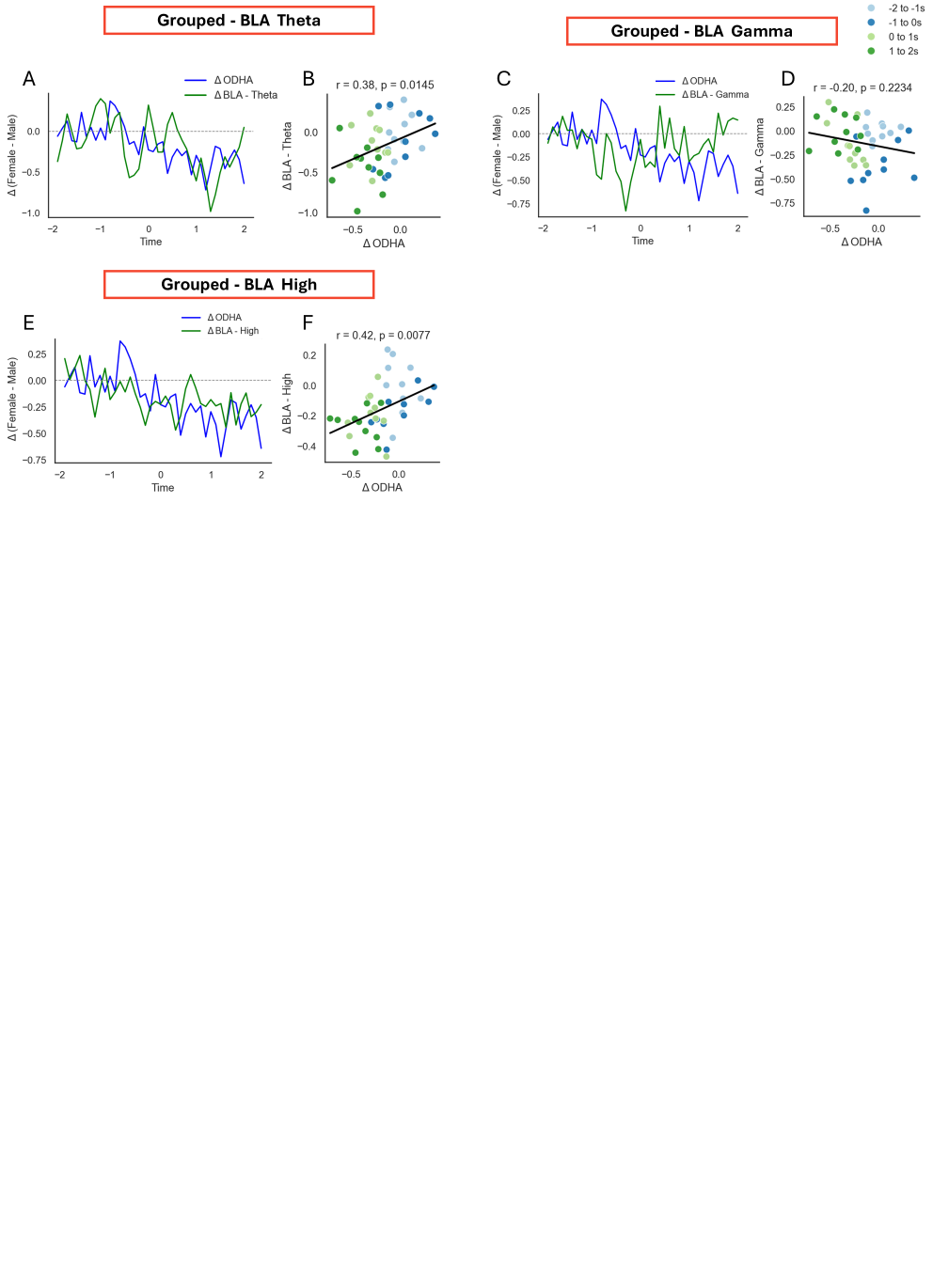

**Figure S9.**

**A.** Time traces of ΔODHA (Z-score Female ODHA minus Z-score Male ODHA, in blue) and ΔBLA LFP theta-frequency power (Z-score Female minus Male, in green) aligned to the onset of approach to the grouped stimulus.
**B.** Scatter plot showing the relationship between ΔODHA (x-axis) and ΔBLA LFP theta-frequency power (y-axis) during investigation bouts with the grouped stimulus. Each point represents a time point, with color indicating time within the event. Regression lines show the fitted linear model. The correlation was assessed using Pearson’s correlation coefficient, and significance was evaluated via permutation testing using 10,000 shuffles. Pearson correlation: r = 0.38, *p* = 0.014.
**C.** Same as (A), but for LFP gamma power.
**D.** Same as (B), but for LFP gamma power.
**E.** Same as (A), but for LFP high-frequency power. **F.** Same as (B), but for LFP high-frequency power. Pearson correlation: r = 0.42, *p* = 0.008.
